# Designing ecologically-optimised vaccines using population genomics

**DOI:** 10.1101/672733

**Authors:** Caroline Colijn, Jukka Corander, Nicholas J. Croucher

**Affiliations:** Department of Mathematics, Simon Fraser University, Burnaby, B.C., V5A 1S6, Canada; Department of Mathematics, Imperial College London, London, SW7 2RH, UK; Department of Biostatistics, University of Oslo, 0372 Oslo, Norway; Helsinki Institute of Information Technology, Department of Mathematics and Statistics, University of Helsinki, 00014 Helsinki, Finland; Parasites & Microbes, Wellcome Sanger Institute, Wellcome Genome Campus, Cambridge, CB10 1SA, UK; MRC Centre for Global Infectious Disease Analysis, Department of Infectious Disease Epidemiology, Imperial College London, London, W2 1PG, UK

## Abstract

*Streptococcus pneumoniae* (the pneumococcus) is a common nasopharyngeal commensal capable of infecting normally sterile anatomical sites, resulting in invasive pneumococcal disease (IPD). Effective vaccines preventing IPD exist, but each of the antigens they contain typically induces protective immunity against only one of the approximately 100 pneumococcal serotypes, which are differentiated by immunogenically-distinct polysaccharide capsules. Serotypes vary in their propensity to cause IPD, quantified as their invasiveness. Vaccines are designed to include serotypes commonly isolated from IPD, but the immunity they induce is sufficiently strong to also eliminate vaccine serotypes from carriage. This enables their replacement by non-vaccine serotypes in the nasopharynx. The emergence of invasive non-vaccine serotypes has undermined some vaccination programmes’ benefits. Recent advances in genomics and modeling have enabled forecasting of which non-vaccine serotypes will be successful post-vaccination. Here, we demonstrate that vaccines optimised using this framework can minimise IPD and antibiotic-resistant disease more effectively than existing formulations in the model, through mitigating the consequences of serotype replacement. The simulations also demonstrate that tailoring vaccines to the pre-vaccine bacterial population is likely to have a substantial impact on reducing IPD, highlighting the importance of epidemiological data, genomics and ecological models as tools for vaccine design and evaluation.

---

Asymptomatic carriage of *S. pneumoniae* peaks in the first five years of life, reaching levels of 25-50% in high-income countries, and 20-90% in low- and middle-income countries^1^. Such high prevalences mean that *S. pneumoniae* strains frequently compete through multiple mechanisms, either during co-colonisation^2^, or indirectly through immune-mediated interactions^3^. The polysaccharide conjugate vaccines (PCVs) routinely administered to infants to limit IPD induce strong mucosal immunity to a limited number of serotypes, preventing their carriage, and alleviating some competition for hosts between the remaining broad diversity of circulating serotypes^4, 5^. This results in a serotype replacement process that typically eliminates vaccine types without any reduction in the overall *S. pneumoniae* carriage prevalence^6, 7^. However, PCVs have substantially reduced infant disease just through altering the carried bacterial population, because serotypes differ in their invasiveness: the rate at which they progress from carriage to cause IPD.

Transmission dynamic modelling of the serotype replacement process has made it possible to quantify the competition between vaccine and non-vaccine serotypes^8, 9^. However, understanding which *S. pneumoniae* serotypes will succeed following alterations to the web of competitive interactions remains difficult. Recent population genomic studies have enabled analyses to move beyond serotypes to consider all variable, or accessory, genetic loci^10, 11^. Corander *et al* observed that accessory loci were preserved at “equilbrium frequencies” both between different global locations with different strain compositions, and between pre- and post-vaccination populations^12^. They hypothesised that multi-locus negative frequency-dependent selection (NFDS) explained these observations, based on functional annotation of the accessory genome^12^. Similar models based on multi-locus NFDS have also proved informative when applied to changing *Escherichia coli* epidemiology^13^, and when reformulated to identify strains likely to invade a vaccine-disrupted population^14^.

The first pneumococcal PCV contained seven serotypes (4, 6B, 9V, 14, 18C, 19F, and 23F) selected to minimise the infant IPD burden, based on epidemiological data primarily from North America and Europe^15^. It was also hoped that PCV7 would reduce the proportion of IPD that was antibiotic resistant, a phenotype strongly associated with some of these vaccine serotypes^16^. Although serotype replacement was not a priority concern, as it was not known whether PCV7 would protect against carriage of vaccine types^1, 15^, PCV7 substantially decreased the burden of infant IPD in many countries^4,17,18^ through its effects on the carried pneumococcal population. The consequent post-vaccine serotype replacement seen in IPD isolates resulted in PCV7 being replaced by PCV10 (which expands PCV7 to include serotypes 1, 5 and 7F) and PCV13 (which adds 3, 6A and 19A to those in PCV10)^1^. These expanded formulations are now administered to millions of children across hundreds of countries^1^. However, post-PCV replacement disease in infants remains a problem, with penicillin-resistant meningitis rising in France post-PCV13^19^. More broadly, there has been little overall effect on the proportion of IPD caused by *S. pneumoniae* resistant to commonly-used antibiotics^20, 21^. Further expansion of PCV valency to tackle these issues is limited by the complexity of manufacturing PCVs, as they are among the most expensive vaccines available^22^, with a full course of immunisations costing over $540 per child in the USA^23^.

Older adults also suffer high incidences of IPD, but do not carry *S. pneumoniae* at the high levels observed in children^1^. Hence infant PCV vaccination programmes alter the serotype profile of adult disease through herd immunity^17^. Yet adult and infant IPD differ in their serotype composition, which appears to reflect their invasiveness varying with host age^17, 24^. Hence the focus on reducing infant IPD results in trade offs with decreasing adult IPD. This is particularly apparent in the UK, where there has been a 4% increase in adult IPD post-PCV13^25^. This highlights the risks attendant to reshaping the bacterial population through PCV-associated strain replacement, as the post-vaccine population can have an increased propensity to cause IPD relative to that preceding the immunisation campaign.

Thus there is a tremendous opportunity to design improved PCV vaccines: vaccination is highly effective at shifting the serotype composition of pneumococcal populations, but is undermined by serotype replacement^17^ and incurs high costs^22^. Here, we use the multi-locus NFDS ecological model^12^ and genomic data describing the circulating carriage genotypes^10, 11^ to predict the serotype distributions resulting from hypothetical vaccine designs. This enables the use of optimization to identify vaccine formulations that should suppress invasive vaccine serotypes and prevent replacement by invasive non-vaccine serotypes. We apply this approach to two different settings to both explore the universal principles of partial coverage vaccine design, and propose formulations that we predict will outperform current vaccines in each location by mitigating the effects of replacement.

## Results

### Incorporating ecology into vaccine design

The two datasets to which the modelling approach was applied had distinct circulating serotypes and genotypes (Fig. S1). The first was from Massachusetts, consisting of 616 genomes sampled from nasopharyngeal carriage in children over three winters following the introduction of PCV7, representing a typical Western *S. pneumoniae* population^10, 12^. The second was from the Maela refugee camp on the Thailand-Myanmar border, comprising 2,336 genomes from an unvaccinated population^12, 26^. Based on the number of detected serotypes, approximately 3.47×10^9^ and 1.05×10^13^ 13-valent PCVs are possible in each, respectively. These datasets were used to calculate the population-wide equilibrium frequencies of the accessory loci, as required for the NFDS modelling, and define the simulated genotypes by their serotype, antibiotic resistance phenotype, and the subset of accessory loci they encoded (Materials and Methods). To enable computationally efficient simulation of how these populations are forecast to respond to arbitary vaccine designs, the multi-locus NFDS model of *S. pneumoniae* ecology was reimplemented in a deterministic form using ordinary differential equations^12^ (Materials and Methods). This version successfully replicated the restructuring of pneumococcal populations following vaccination (Fig. S2). The simulated dynamics are initially driven by vaccination perturbing the population through imposing a fitness cost on those serotypes included in the proposed formulation, followed by a return to an equilibrium under NFDS. Yet the same formulation drives different post-vaccine outcomes in the two locations, due to the different genotype compositions of the carried populations.

These changes in population composition typically stabilised after a decade (Fig. S3), in agreement with epidemiological data^27^. We therefore employed Bayesian optimisation and genetic algorithms to select hypothetical vaccine formulations, and evaluated their impact on IPD 10 years post-vaccination (Fig. S4), using one of three distinct criteria: (1) low infant IPD, (2) low overall infant and adult IPD or (3) low overall antibiotic-resistant IPD.

As NFDS modelling only simulates the carried population dynamics, calculating the IPD burdens used for optimisation requires estimating serotypes’ invasiveness. Invasiveness was separately estimated for infants and adults, as IPD in the two age groups has different serotype compositions^24, 25^, despite adult herd immunity from infant-only vaccination programmes indicating that both emerge from the same carriage population^16, 24^. Hence a meta-analysis of matched carriage and IPD serotype surveys was used to calculate the odds ratios for a serotype being isolated from IPD relative to carriage, relative to all other serotypes detected in the population (Materials and Methods, Table S1, S2). We found that invasiveness odds ratios were broadly similar in adults and infants, with the epidemic serotypes (1, 5, 7F and 12F) more invasive than the paediatric serotypes (6A, 6B, 19F and 23F; Fig. 1A)^16^. Several (8, 12B, 13, 9L, 9N, 20 and 29) had a relatively elevated propensity to cause disease in adults, but little evidence was found of serotypes being highly invasive only in infants (Fig. S5, S6).

**Fig. 1.**
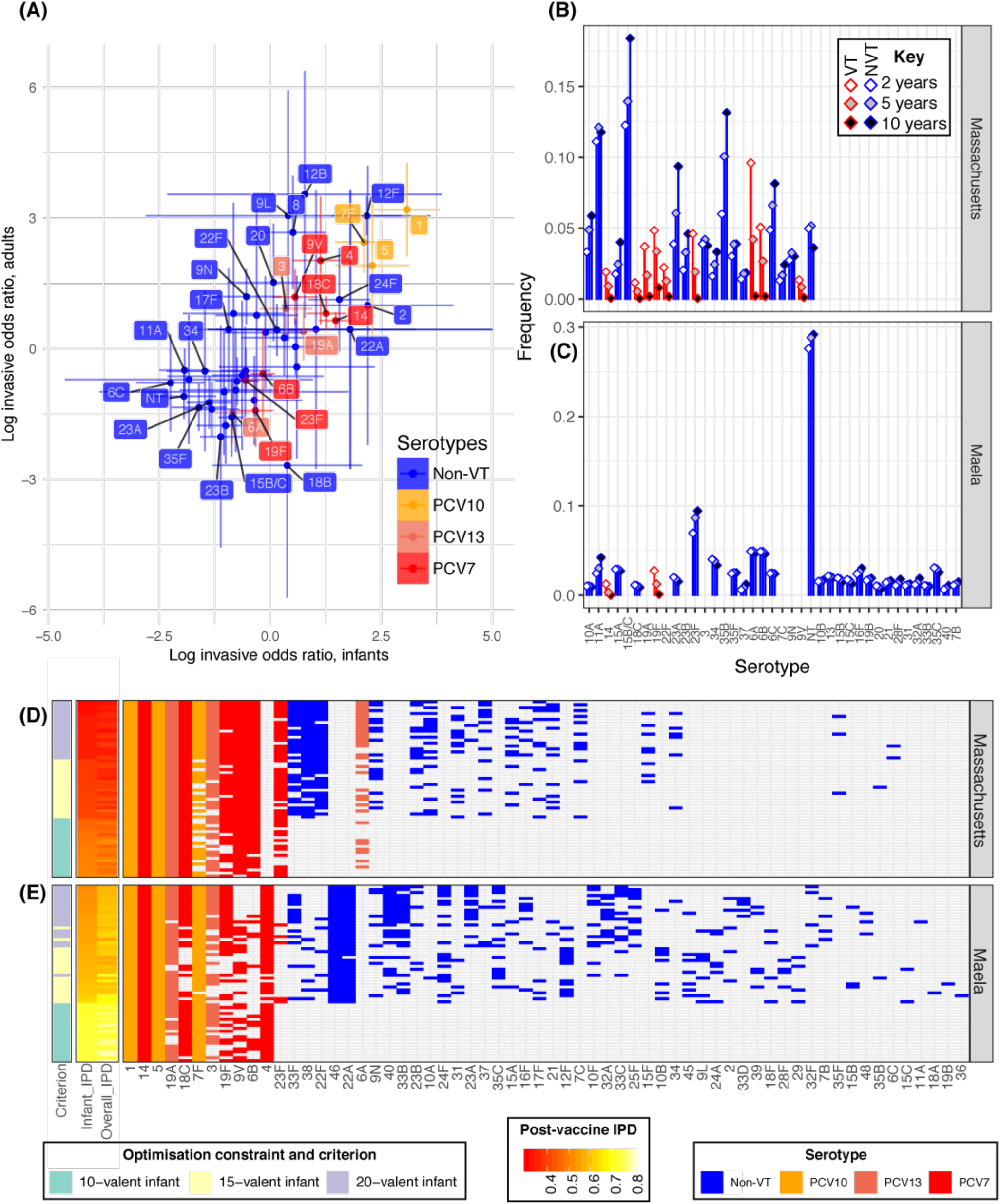
Optimising conjugate vaccines to minimise disease in different demographics (**A**) Invasiveness odds ratios for calculated for pneumococcal serotypes in infants (defined as being under five years old) and adults (all older ages). Points and 95% confidence intervals are plotted on a logarithmic scale and coloured according to the licensed vaccine in which they are found, if any. (**B & C**) Predicted changes in serotype frequencies following introduction of vaccine formulations found to be optimal (among 15- valent vaccines) for minimising infant invasiveness, in (**B**) Massachusetts and (**C**) Maela. (**D & E**) Heatmap summarising the PCV formulations identified optimising for minimising infant IPD under different constraints in (**D**) Massachusetts and (**E**) Maela. The first column shows the constraint on optimisation (15-, 20- or 7-valent vaccine); the adjacent heatmaps show the predicted level of IPD in infants (by which the rows are ordered), and the overall population; and the grid shows the composition of the vaccines, with included serotypes indicated by cells coloured according to their presence in licensed vaccines.

### Minimising infant IPD

We first designed PCV formulations to minimise infant IPD. Optimisation was run with one of three different constraints: maximum valency of 15; maximum valency of 20; or a maximum valency of 10, limited to the constituents of PCV13. These latter formulations are known to be feasible, as the constituent antigens already feature in vaccines, and their cost would be below that of PCV13. We constrained the formulations to include serotypes 1, 5 and 14, which are rare in carriage but highly invasive, and mandatory antigens for a vaccine to be eligible for subsidised introduction into lower-income countries^1^.

In Maela, both 15- and 20-valent PCV formulations were predicted to lower infant IPD to a substantially greater extent than PCV13 (Fig. 1B, C). The best-performing vaccines were those containing highly invasive serotypes, including both serotypes found in current PCVs (e.g. 4, 7F, 18C 19A) and those not yet included in licensed formulations (e.g. 22A, 24F, 46). The other included serotypes differed with the PCV design valency; 10B and 12F were often present in successful 15-valent formulations, whereas better-performing 20-valent formulations often contained 23A, 33F, 33B and 40. Even formulations containing only a subset of the PCV13 components could outperform the 13-valent vaccine, for example by omitting low-invasiveness serotypes 6A, 6B and 23F. Retaining these serotypes in the carried population prevents them from being replaced by higher-invasiveness alternatives following vaccination.

In the Massachusetts dataset, 15-valent formulations could only slightly outperform PCV13 in terms of forecasted infant IPD, whereas 20-valent formulations were more consistently superior. The most frequently added non-PCV13 serotypes were the moderately common and invasive 22F, 33F and 38, resulting in populations dominated by low-invasiveness serotypes (Fig. S7, S8). Surveillance of infant IPD after introduction of higher-valency PCVs has identified 22F as the most common causal serotype, with 33F and 38 also problematic^28^, suggesting that these formulations are likely to perform well in many settings. Similarly, serotypes 12F and 24F have substantially increased in infant IPD incidence following PCV13 administration in the UK and France, respectively^19, 25^. Furthermore, the formulations we identified are predicted to substantially out-perform PCVs composed simply of the serotypes contributing the highest burden of IPD in the starting population (Fig. S9).

### Reducing population-wide and antibiotic-resistant IPD

We then optimised to identify vaccines that would minimize combined infant and adult IPD, with a 50% weighting on each (Fig. 2A). The resulting formulations in Massachusetts do not include serotype 6A, unlike the PCVs designed to minimise infant IPD only (Fig. 1D), likely due to the risk of replacement by serotype 6C, which has a high invasiveness in adults. Indeed, 6C was first identified following the introduction of PCV7 because of its high propensity to cause adult IPD^29^. Similarly in the Maela data, vaccines producing a lower overall infant and adult IPD burden do not contain 19F, unlike those vaccines optimised for minimising infant IPD, likely reflecting the risk of replacement by serotypes with higher invasiveness in adults. PCVs minimising overall IPD instead commonly feature serotype 9N in both locations, which could have helped avoid the substantial increases in adult IPD with serotype 9N recorded in the UK and Sweden post-PCV13^25, 28^. Also frequently included in optimal formulations were 6C, in Massachusetts, and 13, in Maela, reflecting the elevated invasiveness of these serotypes in adults. IPD has a higher mortality rate when the pathogen is resistant to antibiotics^30, 31^. Vaccines can indirectly reduce antimicrobial-resistant (AMR) disease through lowering antibiotic consumption by limiting bacterial disease^32^. Here, we optimise PCVs to directly reduce AMR disease in the absence of any change in antibiotic consumption. The model assumes resistance loci are maintained at their equilibrium frequencies in the carriage population by NFDS^12, 33^, likely driven by levels of antibiotic consumption^34^; however, no such assumption applies to the set of isolates causing IPD. Therefore we optimised vaccine formulations to minimise AMR IPD across infants and adults. We defined a score for each genotype according to the number of loci it contained that were associated with resistance to penicillins, macrolides, co-trimoxazole and tetracyclines, and we penalised strains with resistance to multiple classes of antimicrobials (Materials & Methods). The score’s distribution was highly heterogeneous across both populations, with most serotypes pansusceptible, but a few associated with high levels of multidrug-resistance (Fig. 3A); the paediatric serotypes 19F and 23F, along with 9V and 19A, were associated with resistance in both populations, but the distribution was otherwise quite dissimilar (Fig. S10). We found that vaccine formulations that minimised highly-resistant IPD after vaccination contained serotypes 9V, 19A, 19F and 23F in both populations: 6A and 15A were additionally included in formulations for Massachusetts, where they were associated with AMR^10^. The designed formulations facilitate the success of pan-susceptible serotypes (11A in Massachusetts; 6A and 11A in Maela), low-invasiveness AMR serotypes (6C, 23A and 35B in Massachusetts; 6A and unencapsulated non-typeables, or NTs, in Maela) and isolates only resistant to second-line treatments (e.g. some 15B/C in Massachusetts). While the distinct objectives of minimising overall and AMR IPD require an inevitable trade-off, it was small (Fig S11).

**Fig. 2.**
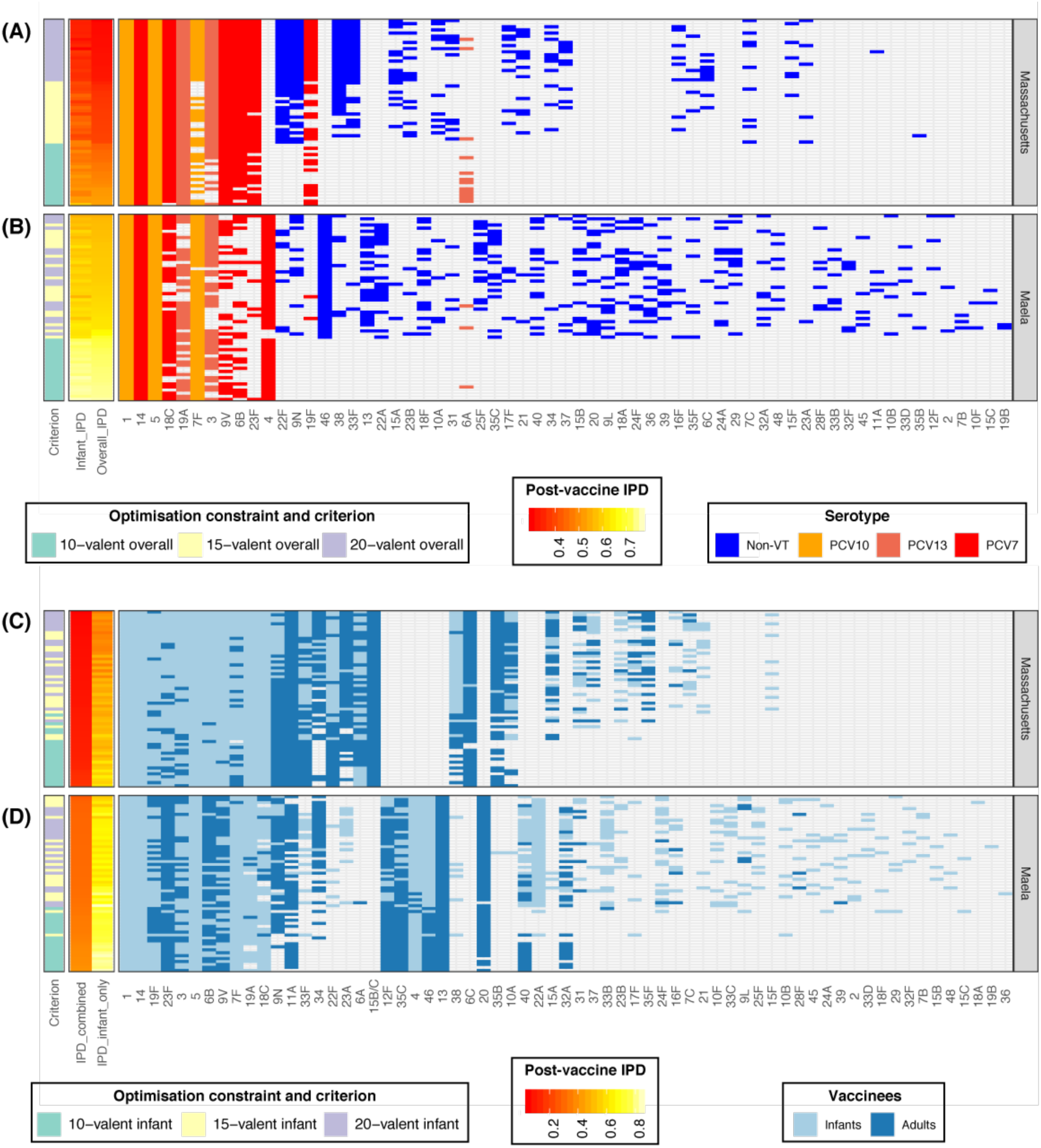
Vaccine strategies for minimising population-wide IPD. **(A & B)** These heatmaps summarise the infant-administered PCV formulations identified optimising for minimising both infant and adult IPD under different constraints in (**A**) Massachusetts and (**B**) Maela, as described for Fig. 1D,E, except that the rows are ordered by the predicted post-vaccination overall IPD burden. This assumes herd immunity induced by the infant vaccination campaign would also eliminate the vaccine serotypes from adult IPD. **(C & D)** Combined strategies in which complementary adult vaccines were designed for each of the infant vaccinations shown in Fig. 1D,E for **(C)** Massachusetts and **(D)** Maela. The complementary adult vaccines provided protection against the 10 serotypes predicted to cause the most disease in adults 10 years after the introduction of the infant-administered vaccine. The adult-administered vaccines were assumed not to drive herd immunity. On each row, the light blue cells define the infant-administered formulation, and the dark blue cells define the adult-administered formulation. These are ordered by the estimated overall IPD level across infants and adults, shown by the IPD heatmaps. Infant-administered vaccines were again assumed to eliminate vaccine serotypes from adult IPD through herd immunity.

**Fig. 3.**
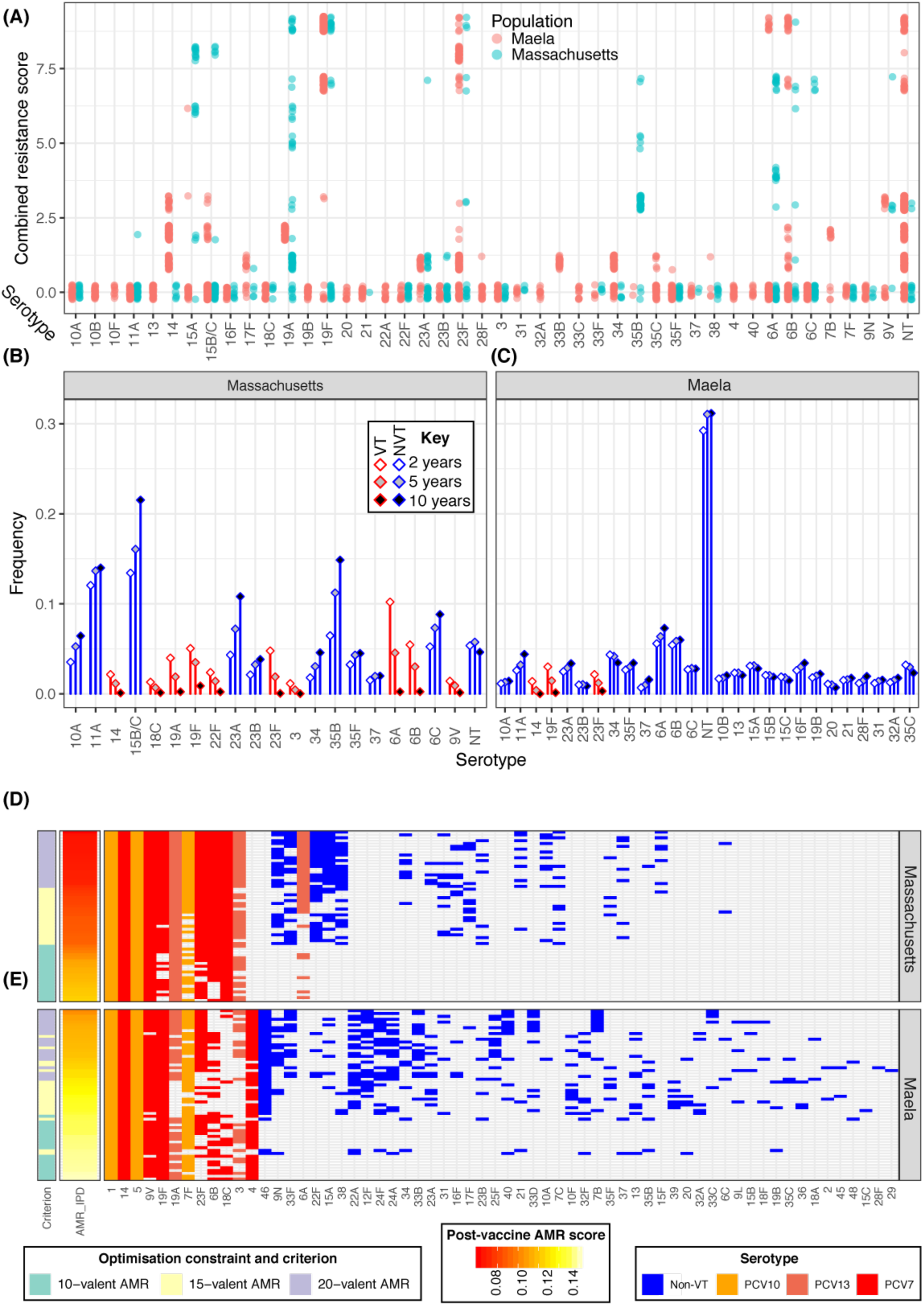
Optimising conjugate vaccines to minimise AMR disease. (**A**) Distribution of AMR score by serotype across the two populations. (**B & C**) Predicted changes in serotype frequency following introduction of 15-valent vaccine formulations found to be optimal for reducing AMR infant IPD in (**B**) Massachusetts and (**C**) Maela, as shown in Fig. 1B,C. (**D & E**) Heatmap summarising the PCV formulations identified optimising for minimising AMR infant IPD under different constraints in (**D**) Massachusetts and (**E**) Maela, as shown in Fig 1D,E.

### Designing protein-based vaccines

In addition to capsular polysaccharides, pneumococci express immunogenic surface proteins. Vaccines based on these proteins have had some success in early-stage trials^1^, but to date these vaccines have typically been based on proteins that are conserved across all isolates^35^. Such formulations cannot exploit intraspecific competition to suppress invasive or resistant variants^8^. However, there are antigenic surface proteins^36, 37^ present in only a subset of pneumococcal isolates (Table S3; Fig. S12), and these could in principle be used to create vaccines targeting a subset of strains, which would re-shape the population in the same manner as PCVs. We modelled antigenic proteins having a vaccine efficacy half that of polysaccharide capsules, and explored vaccine formulations containing combinations of up to twelve intermediate-frequency antigenic proteins (Fig. 4). Resulting successful formulations consistently contained an allele of the zinc metalloprotease ZmpD, which is indeed enriched in invasive serotypes such as 9V and 14 in both populations (Fig. S13, S14). Little other similarity was observed between formulations designed for Massachusetts and Maela, except inclusion of the serotype 9V-associated pilus protein RrgB1. The protein-only vaccine formulations were not predicted to perform as well as PCVs; this may also reflect the higher specificity of PCV targeting, enabling more precise manipulation of the population than is possible with these protein antigens.

**Fig. 4.**
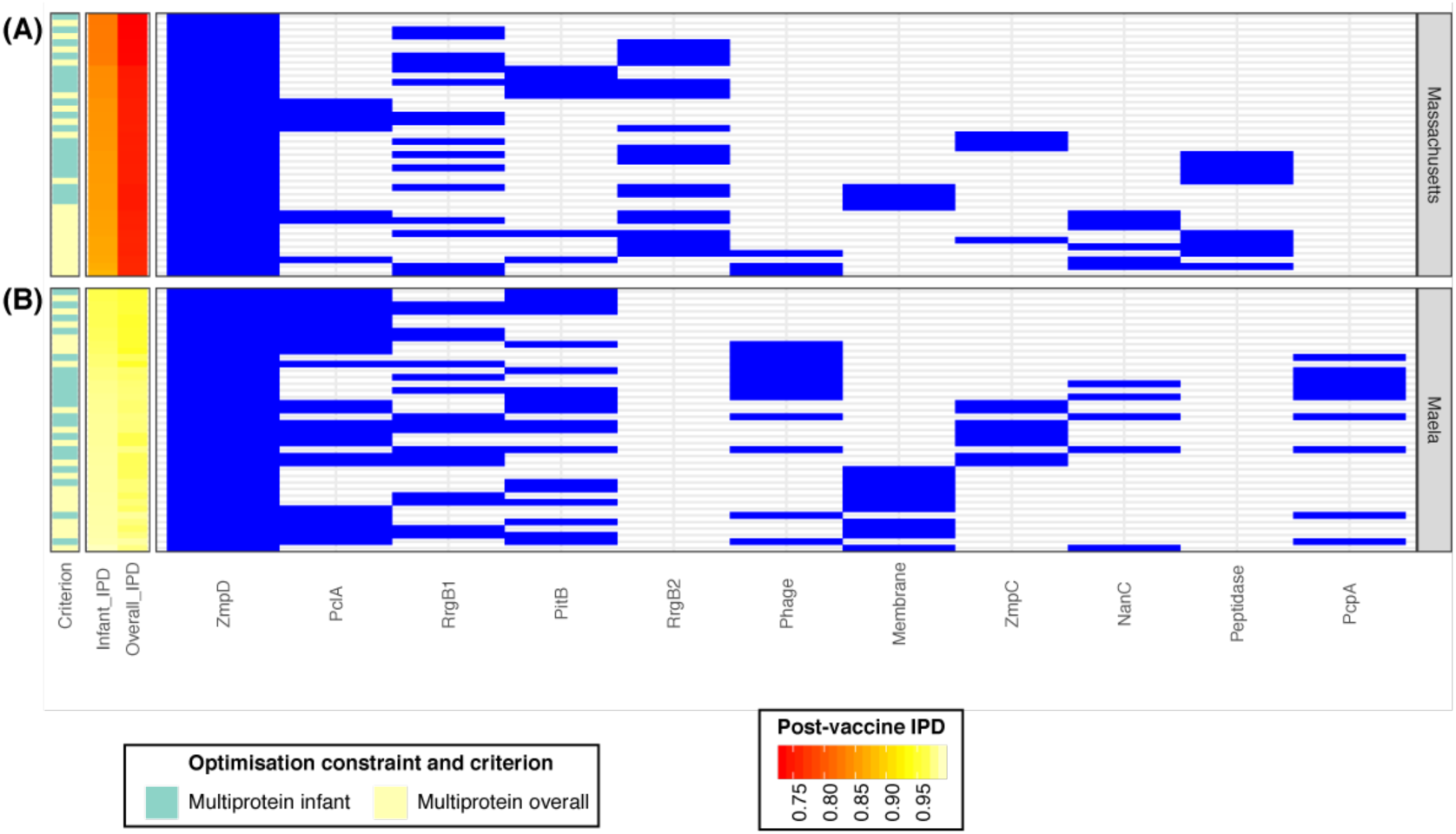
Optimising multiprotein vaccines to minimise infant IPD. The heatmap summarises the protein-based formulations identified when optimising for minimising infant IPD, using an unlimited combination of immunogenic proteins found at intermediate frequencies in the pneumococcal population, in (**A**) Massachusetts and (**B**) Maela. Results are displayed as described for Fig. 1D,E.

Antigenic proteins could also be used as the carrier protein in a PCV; indeed, *Haemophilus influenzae* Protein D is in the currently-licensed PCV10^38^. We optimised to find vaccines based around each one of the twelve variable protein antigens as the carrier, with anti-protein immunity again assumed to be half as effective as anti-capsular immunity (Fig. S15). In Massachusetts, 15-valent formulations with carrier proteins had consistent serotype compositions, and while these occasionally outperformed capsule-only 15-valent PCVs, they were not as effective as 20-valent PCVs. In Maela, some 15-valent formulations with carrier proteins were predicted to outperform capsule-only 15- and 20-valent PCV formulations in both infant and overall IPD. These vaccines’ capsule content depended on the carrier, favouring 22A and 24F (which express few of the carrier protein antigens) over 4, 9V and 19F relative to the carrier-unspecified PCV15 formulations. However, few clear relationships existed between capsules included in the PCV and expression of the carrier protein antigen, nor the frequency of the carrier protein in the population. Therefore, pneumococcal carrier proteins appear promising for PCV design, but their consequences for population structure are hard to predict and likely vary between locations.

### Age-specific vaccine design

Across all criteria, expansion of infant-administered PCV valency was predicted to result in diminishing returns in terms of reducing IPD (Fig. S16). Given serotypes’ differential invasiveness in infants and adults, a more effective strategy may be to develop paired infant-administered and adult-administered vaccines. Currently, many countries offer older adults a 23-valent non-conjugate polysaccharide vaccine that includes all 13 PCV13 serotypes^1^, whereas PCV13 itself was licensed for use in adults in the USA in 2011^39^. However, following the widespread administration of an infant PCV, herd immunity usually suppresses vaccine serotypes across the population, and consequently they contribute little to the IPD burden in unvaccinated adults after approximately seven years post-vaccine introduction^17^. Adult IPD may instead be most efficiently combated by administering a vaccine designed to complement a particular infant PCV, by targeting the serotypes expected to cause adult IPD in the post-vaccine population. The model’s ability to forecast the post-PCV bacterial population, combined with measures of serotypes’ invasiveness in adults, makes it possible to identify which serotypes are predicted to cause the most adult IPD following the establishment of herd immunity by an infant-administered PCV. As adults are not thought to contribute substantially to pneumococcal transmission, such a ‘complementary adult vaccine’ (CAV) would not be expected to reshape the carried bacterial population. We designed such CAVs to complement our optimized infant vaccine formulations, choosing the 10 serotypes that contributed most to adult IPD in the model 10 years after the infant-administered vaccine’s introduction (Fig. 2, S17, S18). Assuming a 90% reduction in invasiveness for CAV serotypes, adult vaccination overcomes the diminishing returns of infant PCV expansion, typically reducing overall IPD by ∼50% relative to an infant-administered vaccination only strategy (Fig. 5A,B). CAVs tended to include both serotypes with elevated invasiveness in adults (6C in Massachusetts; 3, 9N and 12F in Maela), and low-invasiveness serotypes common in the post-infant vaccination population (11A and 34 in both; 15B/C, 22F, 35B in Massachusetts; 13, 20, 23F and 35C in Maela). Many of these (e.g. 6C, 13, 34, 35B, 35C, 35F and 40) do not feature in any currently-available vaccine administered to adults^1^.

**Fig. 5.**
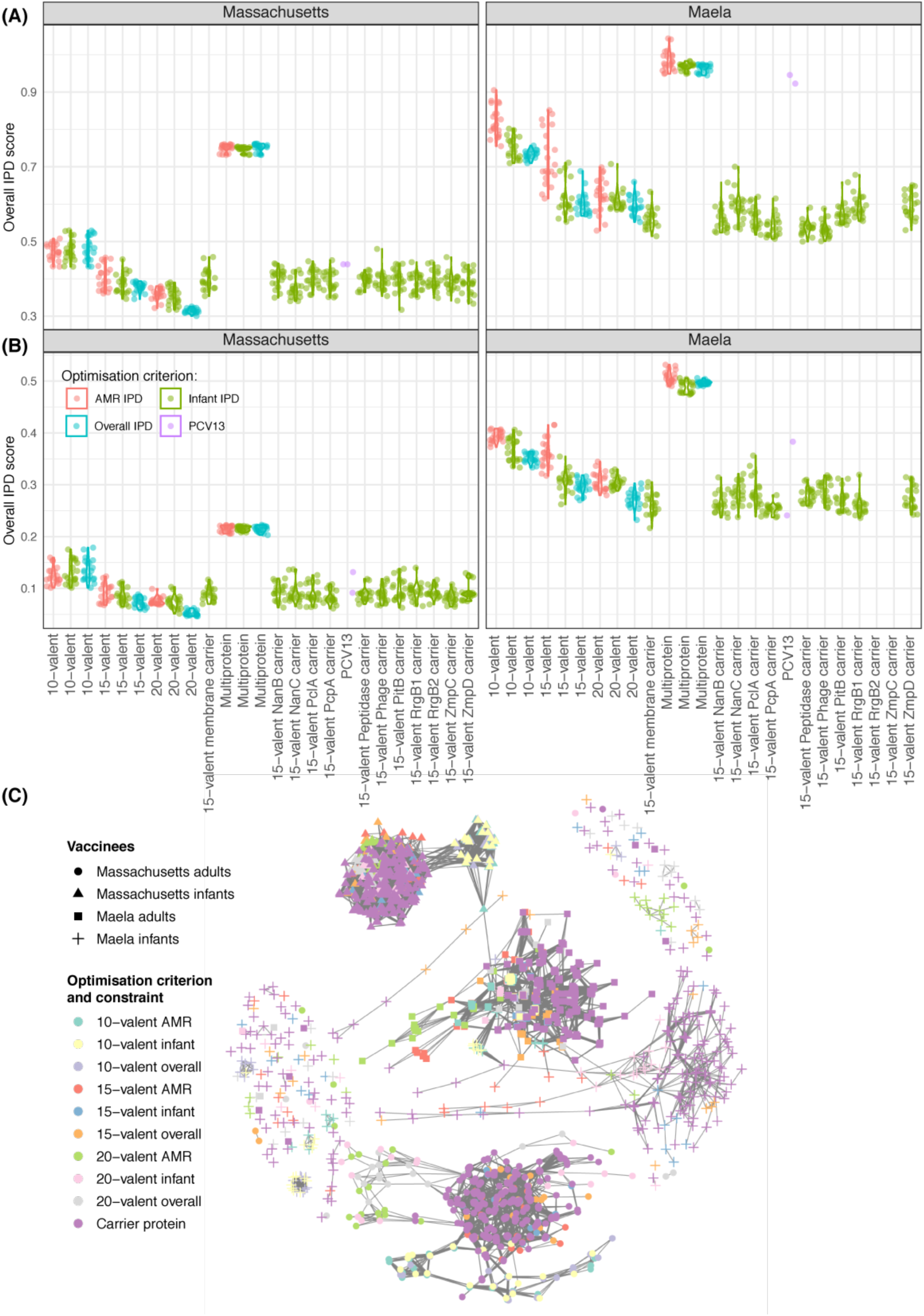
Summarising the effectiveness of different vaccination strategies. **(A)** Violin plots showing the predicted overall IPD burden 10 years post-vaccination in Massachusetts and Maela for all optimal infant-administered vaccine formulations identified in this work. Points are coloured according to the criterion for which they were optimised, with purple points representing corresponding estimates for PCV13. **(B)** Violin plots showing the same estimates with the introduction of CAVs appropriate to each infant-administered vaccine. **(C)** Network summarising the optimal vaccine formulations identified in this work. Each node corresponds to a vaccine formulation, with its colour reflecting the optimisation constraint and criterion, and its shape indicating the intended recipient population. Edges link similar vaccine formulations, identified by applying an empirically-determined threshold to the distribution of pairwise Jaccard distances (Fig. S20).

## Discussion

In many pathogens, interventions are designed using models that do not feature genomic data and the ecological forces driving population dynamics, or using genomic data as a static representation of a pathogen population. However, transmission-blocking vaccines and treatments are continually undermined by pathogen evolution. Our work shows how integrating genomics and modelling can provide new ways to address this major problem. This analysis identified a set of pneumococcal vaccines, each of which was designed to be highly effective for a defined starting population, a design constraint, and an optimisation criterion specifying the type of IPD to be minimised. For each of the infant-administered vaccines expected to alter the carried population, we defined complementary adult-administered vaccines to further reduce the population-wide burden of IPD. These age-specific vaccines can therefore be designed to maximally benefit the respective vaccinee demographics, thereby avoiding one generation enduring elevated risks in order to benefit another.

We illustrate the relationships among the high-performing vaccine formulations with a network in which two formulations are linked if they share a threshold Jaccard similarity level (Fig. 5C, S20). There are four main groupings, corresponding to infant- and adult-admininstered vaccines in the two populations. For each of these four groups, we employed logic regression^40^ supported by manual refinement to summarize the optimal PCV formulations (Table S4). The core specification for infant-administered PCVs for Massachusetts-like populations includes 18C and 19A, which were present in high-performing designs optimised for infant or overall IPD; other effective formulations also have 6B or 9V, and at least three of 19F, 6A, 23F, 3, 38, 7F, 33F and 22F (Fig. S19). Complementary adult vaccines instead should have a core of 11A and 15B/C; one of 23A, 6C, 9N and 10A; and one of 35B, 6A and 33F. In the Maela population, highly-performing formulations for infant-administered PCVs contained serotypes 1, 14, 46 and 5; and at least four of 24F, 22A, 40, 4, 10F, 7F, 19A, 18C, 9L, 19F, 35C, 3, 33C, 9V, 23B, 15A, 15B/C, 36, 32A, 45, and 16F. CAVs contained at least four of 23F, 13, 9N, 19F, 35C, 6B, 20, 3, 9V and 34; and at least one of 24A, 21, 40, 13 or 45.

This underscores the finding that an optimal formulation will depend on the circulating bacterial population, and whether it is expected to block transmission in infants, or only prevent disease in adults. We find that customizing vaccines in this way is likely to produce considerable benefit relative to the global use of a single formulation, particularly if costs are reduced in each location through limiting the antigens in the vaccine to those most important for the local bacterial population (Fig. S21). This argues for a focus on broadening the portfolio of licensed formulations, rather than expanding usage of a single formulation. In particular, expanding usage of vaccines designed for Western populations in locations like Maela may be very much sub-optimal and is likely to be very costly; we forecast a post-vaccine infant IPD average odds ratio of 0.88 for PCV13 compared to 0.55 for a 15-valent design from this analysis, and 0.72 for a 10-valent vaccine whose components are already in PCV13. The first plans for country-specific PCVs are currently being implemented in India^1^, although it will be some years before surveillance data can be used to evaluate its impact.

These conclusions are subject to three principle sources of uncertainty. Firstly, bacterial ecology remains incompletely characterised; further evidence of NFDS shaping populations, and more precise characterisation of the selective pressures involved, are necessary to confidently forecast the effects of vaccines. Yet our optimised formulations are similarly effective in the absence of NFDS, suggesting they do not critically depend on this process for their success (Fig. S22); instead, simulations featuring NFDS filter out vaccines that at risk of causing harmful serotype replacement. Secondly, the unknown genetic basis of strains’ invasiveness, whether entirely serotype-determined or not, makes estimating the IPD burden difficult. This is particularly acute for a location such as Maela, where many prevalent serotypes are associated with little epidemiological data on their propensity to cause disease (Fig. S23). These poorly-characterised serotypes may emerge as more global concerns as higher-valency PCVs deplete currently-circulating strains. Thirdly, our modelling of serotype replacement is limited by our understanding of global transmission patterns and strain diversity. International sequencing-focussed research projects, and routine genomic surveillance, will help address all three lacunae. These advances can be integrated through the framework presented here to aid vaccine design and, given local surveillance data, inform policy making at a regional level. Combined with recent advances in manufacturing techniques, there is an emerging opportunity to apply the principles of ‘precision medicine’ to ensure PCVs are maximally effective for everyone.

## Materials and Methods

### Meta-analysis of serotype invasiveness

To identify paired samples of pneumococci from invasive disease (IPD) in infants or adults, relative to the circulating carriage population in infants, PUBMED was searched with the following terms on 5th October 2017:

(case[All Fields] OR disease[All Fields] OR episode[All Fields] OR patient[All Fields]) AND (carriage[All Fields] OR carrier[All Fields] OR nasopharyngeal[All Fields]) AND (invasiveness[All Fields] OR “attack rate”[All Fields] OR “type distribution”[All Fields] OR “serotype distribution”[All Fields] OR “serogroup distribution”[All Fields] OR “invasive capacity”[All Fields] OR “invasiveness ratio”[All Fields] OR “odds ratio”[All Fields] OR “carrier ratio”[All Fields] OR (“invasive isolates”[All Fields] AND “carriage isolates”[All Fields])) AND (“serogroup”[MeSH Terms] OR “serogroup”[All Fields] OR “serotype”[All Fields]) AND (“streptococcus pneumoniae”[MeSH Terms] OR (“streptococcus”[All Fields] AND “pneumoniae”[All Fields]) OR “streptococcus pneumoniae”[All Fields] OR “pneumococcus”[All Fields])

This returned 136 results, the abstracts of which were reviewed to identify those in which data could be extracted for meta-analysis at a serotype-specific level of precision. Thirty-four abstracts were found likely to be appropriate. After reading the papers, six did not contain matched disease and asymptomatic carriage samples, and seven further individual studies were rejected due to bias towards particular serotypes or lack of serotype-level reporting, very high co-colonisation complicating analysis of the carriage sample, difficulties using data when stratified by age, or inability to access the raw data. This left 21 studies with matched systematically-sampled and thoroughly serotyped asymptomatic carriage and disease samples^24,41–60^. Within these, isolates of the rapidly-interconverting serotypes 15B and 15C were combined into a single 15B/C category. Samples were then stratified by age and data of vaccine introduction, generating 23 pairs of infant carriage and infant IPD samples (seven of which were post-PCV introduction), and 7 pairs of infant carriage and primarily adult IPD samples (one of which was post-PCV introduction). Logarithmic invasiveness odds ratios were calculated across datasets by fitting linear mixed-effects models using the metafor package^61^ in R. The studies are listed in Table S1, and the data summarised in Table S2.

When calculating IPD burdens, if an adult invasiveness value was not available for a serotype, its infant invasiveness was used instead. If an infant invasiveness estimate was not available, the lowest invasiveness estimate from within the same serogroup was used, where one was available; otherwise a value associated with a similarly rare serotype with a low invasiveness estimate was selected. The invasiveness of vaccine serotypes was not altered in the post-vaccine period, as the pre- and post-PCV invasiveness odds ratios were not substantially altered for vaccine serotypes relative to non-vaccine serotypes in epidemiological data (Fig. S6).

### Model specification

We approximate the stochastic model of Corander *et al*^12^ with a deterministic set of ordinary differential equations describing the evolution of the pneumococcal population in response to a vaccine strategy. We model the same negative frequency-dependent selection (NFDS), in which each intermediate-frequency locus *l* (present at between 5% and 95% prevalence in the initial population) is assumed to have an equilibrium frequency, *e_l_*. This frequency is calculated from the pre-vaccine population. Vaccine-induced immunity perturbs the population through removal of vaccine-type serotypes, meaning the instantaneous frequency of *l* at time *t* after the vaccine’s introduction, *f_l,t_*, may deviate from *e_l_.* As NFDS means loci are most advantageous when rare, genotypes have a high fitness when they are enriched for loci below their corresponding equilibrium frequencies; their elevated reproductive rate returns the loci frequencies towards *e_l_*. Each isolate is defined by its serotype and its genotype, determined by the intermediate-frequency loci it carries. The genotypes are recorded in a matrix *G* with *G_il_*=1 if strain *i* has *l*, and 0 otherwise.

We model the NFDS with a term 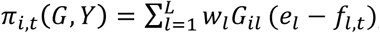, where *L* is the total number of intermediate-frequency loci, and *Y* is a vector whose components *y_i_* are the prevalences of the genotypes, indexed by *i*. The *w_l_* are weights, distinguishing between loci under strong or weak NFDS^12^. The index *i* runs from 1 to *M*, the number of unique intermediate-frequency locus profiles in the model (*M* is 603 for the Massachusetts data and 674 for the Maela data). The NFDS term depends on the prevalence of all the strains in the model because it depends on the frequency of each locus; this couples the strains together. The frequencies are computed from the prevalences:

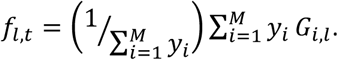

To derive a deterministic model describing the same average population dynamics as ^12^ we use the standard first-order approach, equating the fractional change in a fixed time frame in the two models. This gives 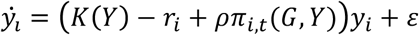, where 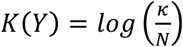 is a term ensuring a carrying capacity of κ (here taken to be 10^5^) and 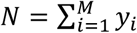. The vaccine strategy is embedded in the *r_i_* which are either a constant *r* (if the serotype of genotype *i* is included in the vaccine), or 0. The constant ρ is the overall strength of NFDS; for the “neutral” simulations exploring robustness to NFDS we set ρ = 0.

The parameters *r* and ρ were fitted to the model of ^12^ to obtain the same rates of decline of vaccine strains and rise in non-vaccine strains following vaccination (Fig. S2), yielding *r* = 0.063 and ρ = 0.165. Equilibrium locus frequencies *e_l_* and weights *w_l_* are as in ^12^.

To reduce the dimensionality in the Maela dataset we model frequencies of clusters of genotypes rather than each individual genotype. We obtain clusters using a graph approach; we create a graph whose nodes are individual genotypes and whose edges join two genotypes if they differ at fewer than 20 loci and have the same serotype and resistance loci. Each of the 674 connected components in this graph is modelled as a genotype; its loci are modelled as those of the component’s highest-degree genotype.

For the data from Massachusetts, the PCV7-associated population dynamics made it important to use the pre-vaccine population frequency of each sequence cluster, taken as a proportion of the carrying capacity, as the initial frequency *y_i_*(0). The Maela samples were collected over a short period in an unvaccinated population, and therefore we model each sequence cluster as equally prevalent initially.

### Optimisation approach

We optimised for three distinct criteria: (1) infant IPD; (2) overall IPD, which equally weighted each serotype’s invasiveness in infants and adults; and (3) AMR IPD, a criterion under which genotypes score highly if they are both resistant and invasive. For a modelled population with prevalences *y_i_*(*t*) of strain *i* at time *t* following the introduction of a vaccine, the infant IPD burden was estimated as 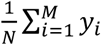 exp (*k_i_*) where *K_i_* is the infant invasiveness log odds ratio of genotype *i*, based on its serotype, and *N* is the total prevalence (which is very close to the carrying capacity). Similarly, the overall IPD burden was calculated as 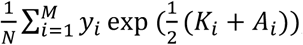, where *A_i_* are the serotype-derived log odds ratios of invasive disease in adults for genotype *i*.

We modelled a resistance score for each isolate and used a logistic model based on minimising the probability of invasive and resistant disease. The score for an isolate is 0 if it is susceptible to penicillin, which corresponds to the isolate having β lactam-susceptible alleles at each of the three relevant penicillin-binding protein-encoding loci ^10, 12^. If the strain appeared to exhibit any β lactam non-susceptibility, this conferred a score equal to the number of loci at which β lactam resistance alleles were present (*n_p_*). If the genotype was also inferred to be macrolide resistant, then *n_m_*(set equal to one) was added to the score; furthermore, if the macrolide-resistant genotype encoded loci conferring resistance to trimethoprim, sulphamethoxazole (the components of co-trimoxazole, cumulatively quantified as *n_c_*), or tetracycline (quantified as *n_t_*), the resistance score was incremented by the appropriate number of resistance loci. In summary, if *I_p_* and *I_m_* are indicators for the presence of any β lactam or macrolide resistance loci, respectively; *n_p_*, *n_m_*, *n_c_* and *n_t_* are the numbers of loci associated with the four described antibiotic classes, the resistance score of genotype *I* is:

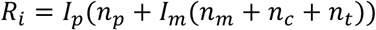

This is broadly motivated by prescribing practices that first use penicillin and, if that is ineffective, a macrolide, followed by less common use of other antibiotic classes. Based on the score, we model a logistic probability^62^ of resistance to treatment as 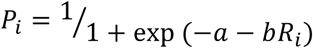 with *a* = −2 and *b* = 0.5. The combined AMR IPD criterion is calculated as 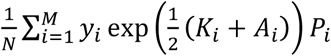, which combines infant and adult invasiveness with the probability of resistance to treatment. We chose to use odds ratios (ORs) rather than log ORs in the criteria because this will drive the optimisations to strongly attempt to suppress highly invasive strains.

Our criteria carry uncertainty because the invasiveness estimates are uncertain. The serotype-based invasiveness in infants and adults (log ORs *K_i_* and *A_i_*) are point estimates with accompanying standard deviations, obtained in the meta-analysis. To assess uncertainty in the criteria, we resampled each serotype’s invasiveness log OR from a normal distribution with mean and standard deviation obtained in the meta-analysis. Each strain was assigned the new log OR corresponding to its serotype, and the criterion was recomputed. Because our criteria feature ORs (not log ORs), the resampled criteria are positively skewed. We illustrate the magnitude of uncertainty in the infant invasiveness estimates in Fig. S23, which shows inter-quartile ranges for the analysis summarised in Fig. 4 of the main text; other criteria are qualitatively similar in uncertainty. We also explored resampling the invasiveness of a serotype in different individual hosts according to the same distribution, reflecting the recognition that the same serotype may have different propensity to cause invasive disease in different individuals. This results in less variance in the objective estimates than is shown in Fig. S5 and S6 because prevalent serotypes’ invasiveness is sampled many times, and the average of these samples is close to the mean (by the central limit theorem); rare serotypes have more variance but as they are rare they contribute less to the objective function.

The model was solved in matlab with the ode15s solver. All prevalences were set to be non-negative, the absolute tolerance was 10^-8^ and the relative tolerance was 10^-5^. Simulating the pneumococcal population over 10 years took between 15 and 30 s (depending on the vaccine strategy). We primarily used Bayesian optimisation in matlab to explore the space of possible vaccine strategies; this is implemented in the ‘bayesopt’ function in the statistics and machine learning toolbox. We constrained the number of serotypes to a 15- or 20-valent formulation, including serotypes 1, 5 and 14, which are mandatory for a PCV to be eligible for subsidised introduction into lower-income countries through the GAVI Advance Market Commitment^1^. We also ‘downsampled’ PCV13, selecting up to 7 of the serotypes in PCV13. The ‘bayesopt’ function uses its own acquisition function to determine where next to search the space of possible strategies; where this failed due to its chosen strategies not meeting our constraints, we used a genetic algorithm (’ga’ in matlab’s Global Optimization Toolbox) with customised mutation and crossover functions to sample vaccine strategies that matched our constraints.

### Model dynamics

We chose to assess the objective functions at a 10-year time point (Fig. S3, S4). While the model has long transient behaviour in the genotype frequencies, this is primarily due to slow drifting amongst very similar genotypes with extremely low rates of change. The objective functions are very similar at the 10, 25 and 50-year time points (Fig. S3).

The equilibria and their stability are not obtainable analytically, even if the logarithmic term were replaced with a polynomial one (e.g. a logistic term, which is a good approximation if the population *N* is near the carrying capacity *K*). In a simplified version of the model in which the population is at this carrying capacity, and in which the migration term is 0, the equilibrium condition can be written 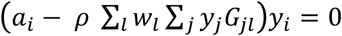, where 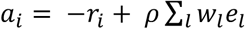 and *e_l_* are the equilibrium locus frequencies. The term 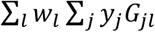 is, in matrix notation, *w^T^G^T^ y*, with *w* the vector of weights *w_l_* and *y* the vector of prevalences *y_i_.* The matrix *w^T^G^T^* has rank 1 (it is a row vector), and a null space of rank *M-1.* This means that if *y\** is a solution to the equilibrium equation such that the term in brackets is 0, then *y*+y_n_* is also an equilibrium solution, for any vector *y_n_* in the null space of *w^T^G^T^*. On this basis we expect that there are many possible equilibria of the system, including also others where for some *i* the term in brackets vanishes and for others the strain is eliminated (so the *y_i_* term in the equilibrium equation vanishes instead). With a polynomial term in place of the logarithmic one, it may be possible to characterize the equilibria using techniques from algebraic geometry to describe the solutions to this high-dimensional polynomial equation. On a practical note, the possibility of multiple equilibria means that the solutions depend on the initial conditions, potentially even after long periods. Hence, carriage data should be used to define the initial conditions as precisely as possible.

### Sensitivity to initial conditions

We resampled the initial conditions of the model in two ways. First, we added Gaussian random noise to the initial prevalence of each genotype, where for each genotype, the standard deviation of the added noise was 10% of the genotype’s starting prevalence. This models the notion that the dataset is correct with regards to which genotypes are present, but uncertain about their precise prevalence. This perturbs the overall IPD burden by less than 1% on average (for example a standard deviation of 0.0027 for an overall IPD burden of 0.41), and a maximum of 2%. We then models the notion that the dataset may not correctly reflect which genotypes are initially present in larger numbers, due to sampling effects. We uniformly chose 10% of the genotypes, and permuted their initial frequencies, thereby allowing some that were not initially modelled as present in higher quantities to be initially present and vice versa. This results in a larger variation than adding 10% noise to all initial conditions (for example a standard deviation of 0.01, compared to an overall IPD burden of 0.41 with the original initial frequencies). Overall the invasiveness objectives remain similarly robust to changes in the initial conditions.

We also resampled the equilibrium locus frequencies, adding Gaussian random noise with a standard deviation of 10% of the default values. The resulting invasiveness varies more than under perturbed initial conditions, which is not surprising given that the specified locus frequencies shape the long-term population dynamics through the frequency-dependent selection term. The resulting invasiveness values had standard deviation of under 5% of the typical objective for the strategy (e.g. 0.018 for an overall IPD burden of 0.41). In the case of this high-performing test strategy with an overall IPD burden of 0.41 (containing serotypes 14, 17F, 18C, 19A, 19F, 22F, 23F, 33F, 38, 6A, 6B, 7F and 9V) the invasiveness ranged from 0.39 to 0.45 under 20 perturbations. The invasiveness criteria all tend to be robust to small perturbations in the locus frequency parameters, and are similarly robust to the locus weights; these have similar effects to perturbations to the equilibrium frequencies.

Figure S22 shows the relationship between formulations’ performance in the model and in the neutral variant in which NFDS does not affect population dynamics.

### Complementary paired formulations

To identify complementary vaccines to minimise adult IPD given an infant-administered PCV that modifies the carried pneumococcal population, we simulated the primary PCV strategy to the 10-year time point. We computed each serotype’s contribution to the total adult IPD burden as 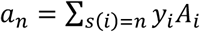, where *s*(*i*) is the serotype of genotype *i*. We included the 10 serotypes making the greatest contributions to adult IPD. To model the updated adult IPD burden we assumed that inclusion in the complementary vaccine would reduce a serotype’s invasiveness in adults by 90%.

### Validating the necessity for machine learning approaches

To test for whether the combination of NFDS modelling and machine learning provides an advantage over a formulation of those serotypes causing the highest burden of IPD, we chose 15-valent formulations that consisted of the top 15 serotypes contributing to infant IPD in the initial population. In the Massachusetts population, the resulting formulation contains types 11A, 14, 15B/C, 18C, 19A, 19F, 22F, 23F, 3, 6A, 6B, 9N and 9V (and in keeping with the rest of the work we would add 1 and 5), and results in an infant IPD burden at 10 years of 0.63. This is higher than both expected following the introduction of PCV13 (0.42), and well above that expected for the best-performing 15-valent strategy we identified in the optimisation (0.37). This strategy contains types 1, 5, 14, 17F, 18C, 19A, 19F, 22F, 23F, 33F, 38, 6A, 6B, 7F, and 9V, allowing it to suppress invasive strains that are not prevalent in the initial data but become so upon elimination of vaccine strains, thereby preventing problematic serotype replacement of the types observed in French infants^19^, UK adults^25^, and elsewhere^17^. Figure S9 shows the predicted serotype distributions under these strategies in the two populations.

## Acknowledgments

We thank Dr. Corinne Levy for sharing epidemiological data.

## Funding

CC was supported by the Engineering and Physical Sciences Research Council of the UK (EP/K026003/1; EP/N014529/1) and by the Government of Canada’s Canada 150 Research Chair program. JC was supported by ERC grant number 742158. NJC was supported by a Sir Henry Dale Fellowship, jointly funded by Wellcome and the Royal Society (104169/Z/14/Z), and by the UK Medical Research Council and Department for International Development (MR/R015600/1).

## Author contributions

CC and NJC designed the study; CC, JC and NJC developed the model; CC and NJC analyzed data; all authors wrote the manuscript.

## Competing interests

CC and NJC have protected the formulations identified in this work. NJC has consulted for Antigen Discovery Inc.

## Data and materials availability

Model code is available at https://github.com/carolinecolijn/optimvaccine

## Supplementary Materials

**Fig. S1.**
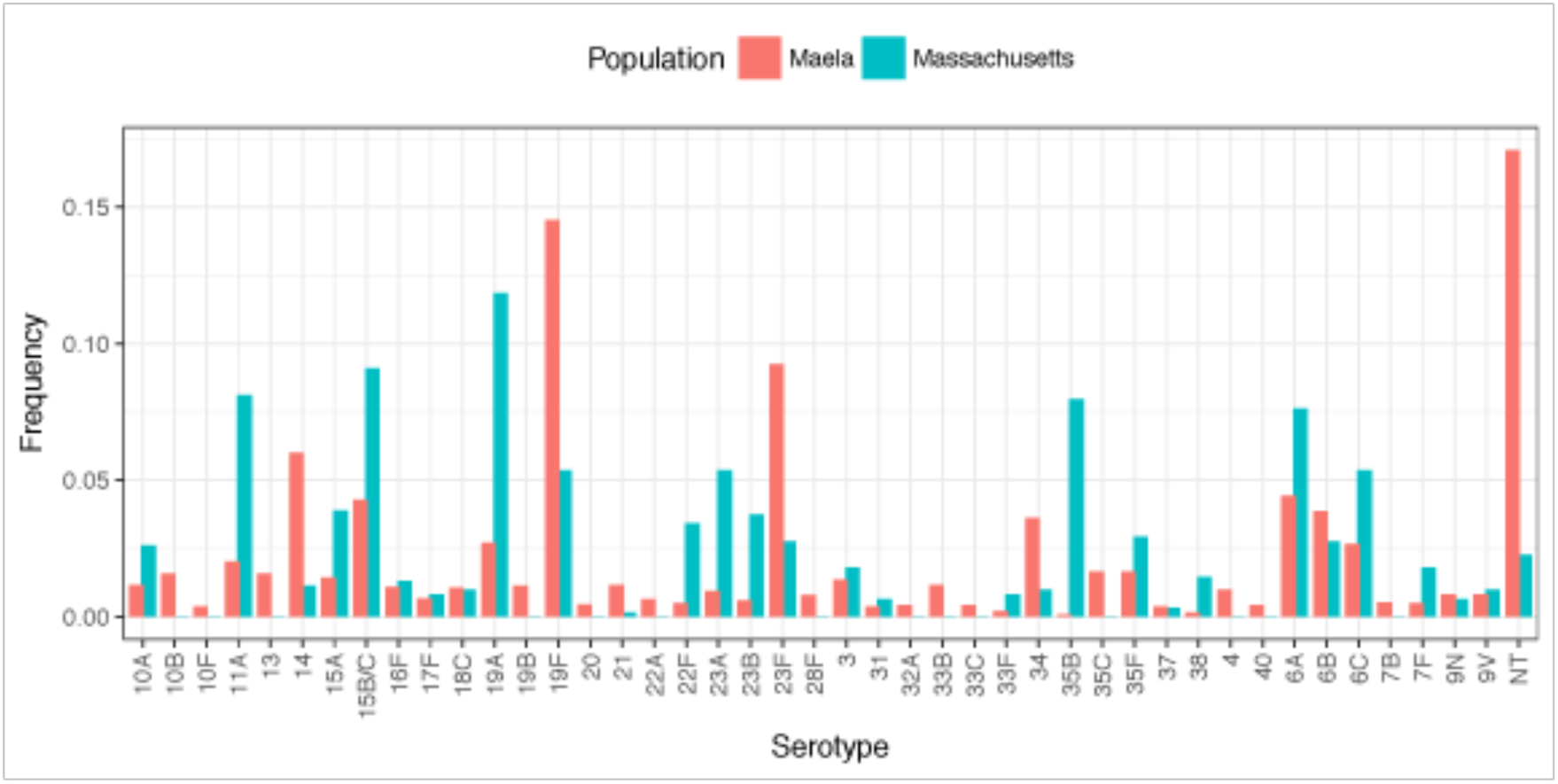
Frequencies of serotypes across the two studied populations; serotypes 15B and 15C, which rapidly interchange but were resolved separately in the Maela dataset, are merged into 15B/C for comparability in this plot.

**Fig. S2.**
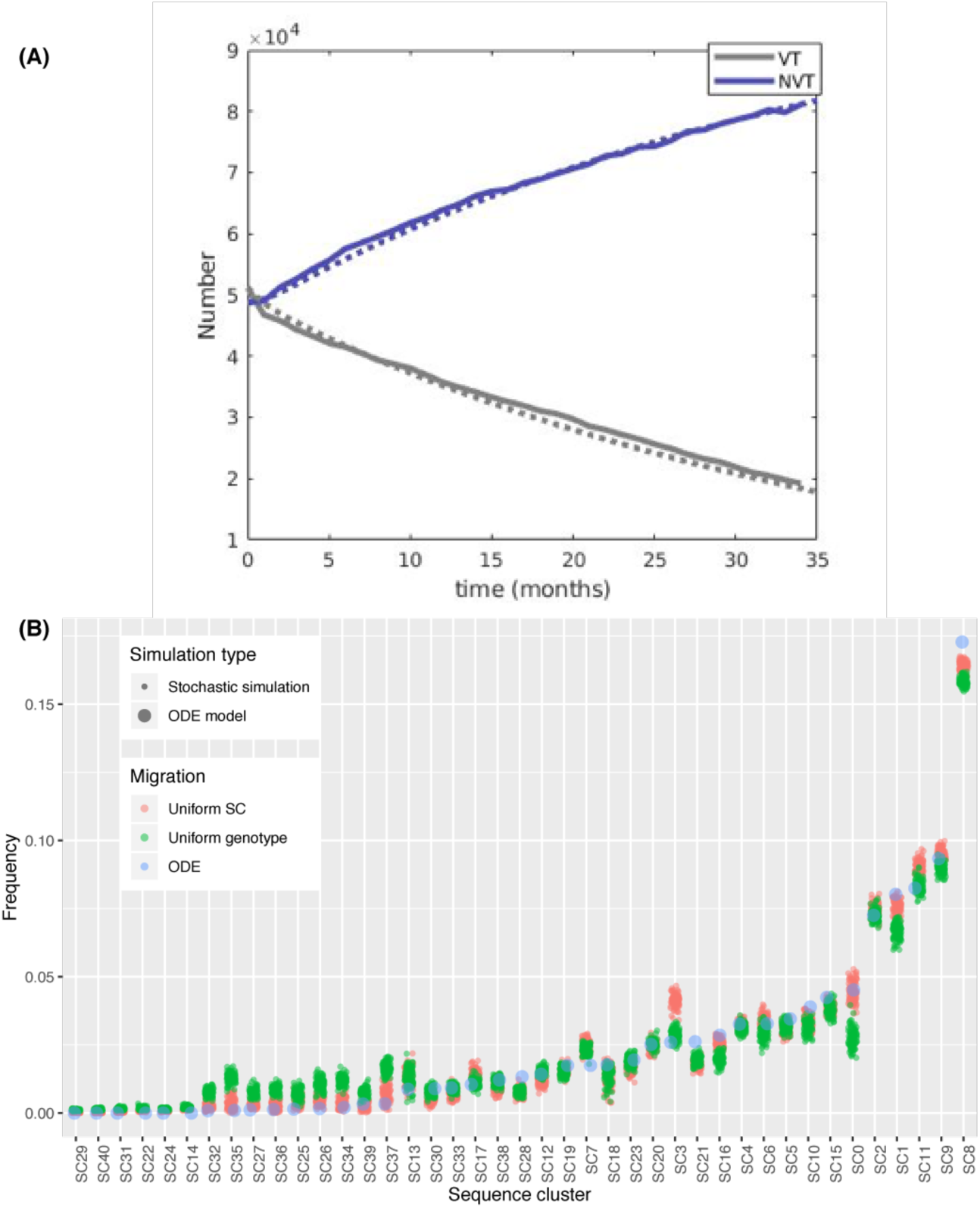
Correspondence between the ODE and stochastic multi-locus NFDS models when simulating the impact of PCV7 on the Massachusetts *S. pneumoniae* population. **(A)** Model fitting. The similarity between the solid lines (stochastic model output) and dashed lines (ODE model output) shows the deterministic ODE model replicates the temporal dynamics of the stochastic version, which was parameterised through fitting to genomic surveillance data. **(B)** Replication of the post-PCV7 population at 10 years. The frequency of sequence clusters, defined in Corander *et al*, in the two model implementations was compared. One hundred replicates of two sets of stochastic model outputs are shown: one set for a uniform migration rate per sequence cluster, which was used to facilitate model fitting, and one set for a uniform migration rate per isolate. These reach slightly different population compositions after ten years. The ODE model necessarily uses a deterministic uniform migration rate per genotype, which is intermediate between the two mechanisms implemented for the stochastic model: each genotype may represent multiple isolates, and each sequence cluster contains multiple genotypes. Appropriately, these simulations arrive at a third equilibrium, in which each SC’s frequency matches that in at least one, and usually both, of the stochastic model outputs. This is consistent with an accurate replication of the NFDS mechanics, and the uncertainty of the migration process, given the current paucity of well-sampled carriage collections from the wider *S. pneumoniae* metapopulation.

**Fig. S3.**
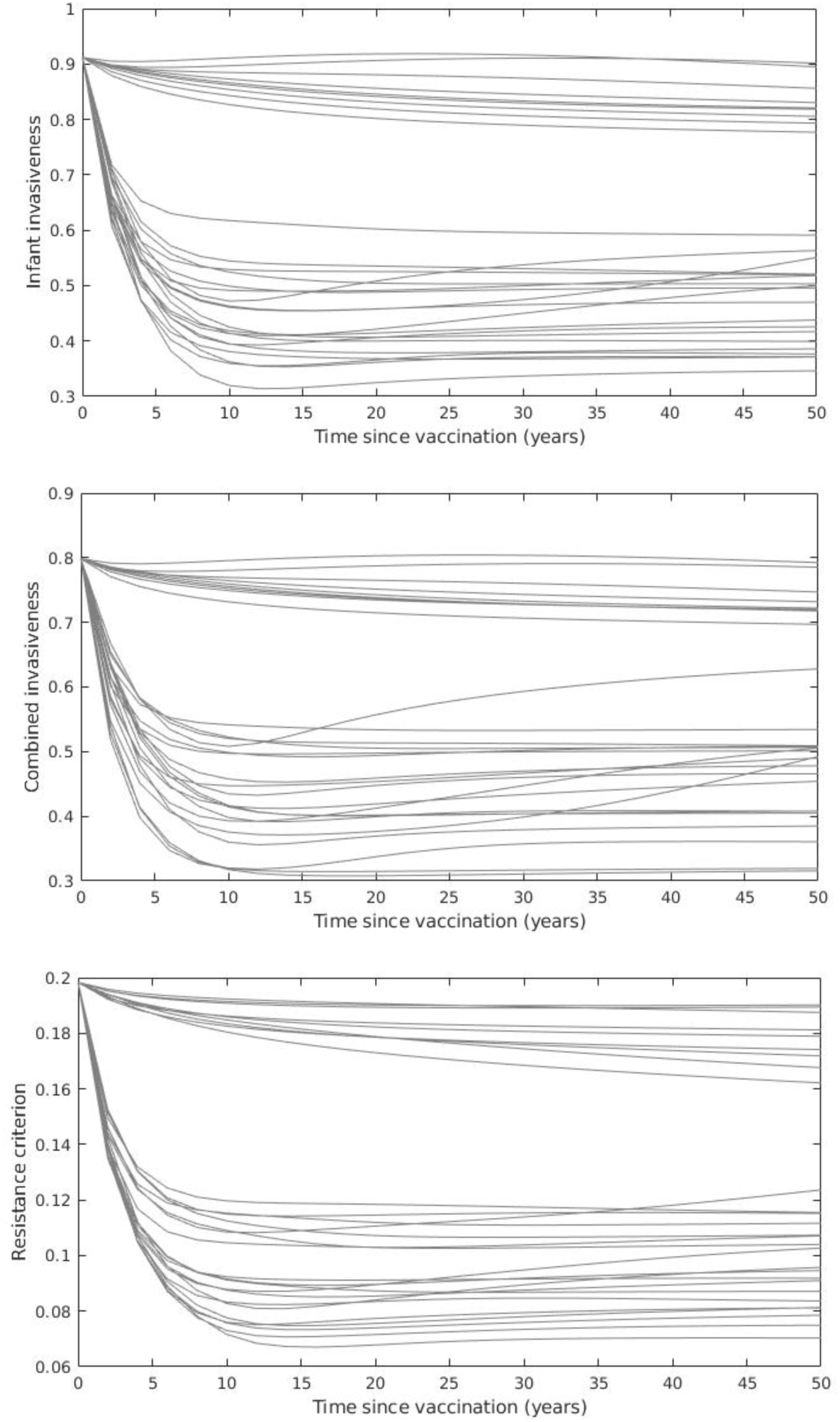
The three measures of IPD burden used to optimise vaccine formulations as a function of time for a random selection of strategies simulated as being implemented in the Massachusetts population. By 10 years the criteria have either reached their stable levels or, in rare cases, reached a minimum from which they slowly rise over the subsequent 40 years. Formulations with low invasiveness at 10 years tend to have correspondingly low values at 25 years, as can be inferred from the lines rarely crossing after 10 years, with slow drift in the few exceptions. We chose to evaluate the criteria at 10 years; in practice we suggest that continued surveillance would enable the development of vaccines that would mitigate longer-term rises in invasiveness or resistance.

**Fig. S4.**
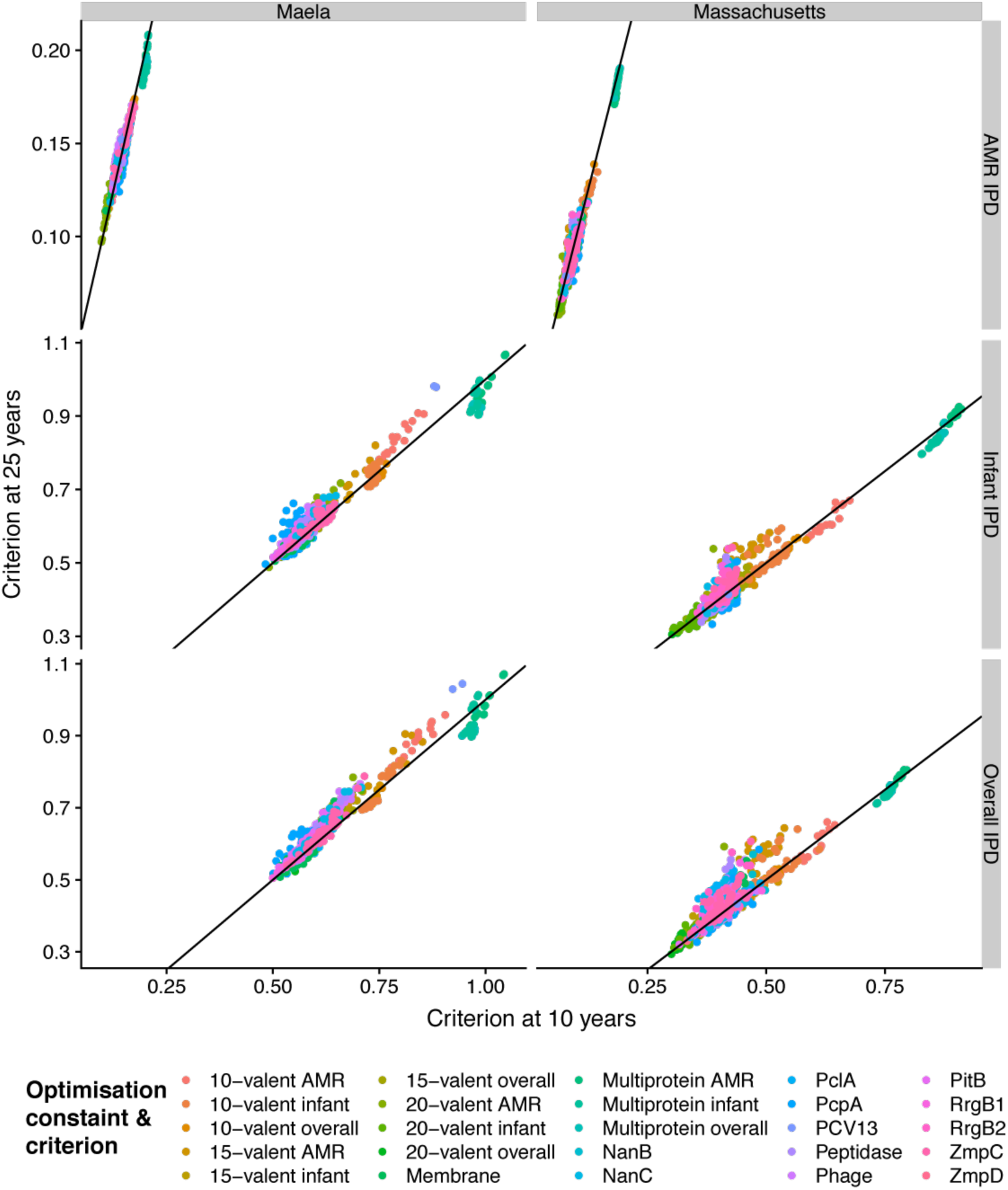
Correlation between IPD burden measures used for vaccine optimisation at 10 and 25 years post-vaccination. Plots are separated by population and IPD burden measure; points are coloured by the constraint on the formulation, and the criterion used for optimisation. The line of identity is marked in black. The IPD measures are strongly correlated at the two timepoints, indicating that while the model dynamics have long transient behaviour driven by drift among similar genotypes, the IPD burden criteria converge towards a feasible-time value relatively early.

**Fig. S5.**
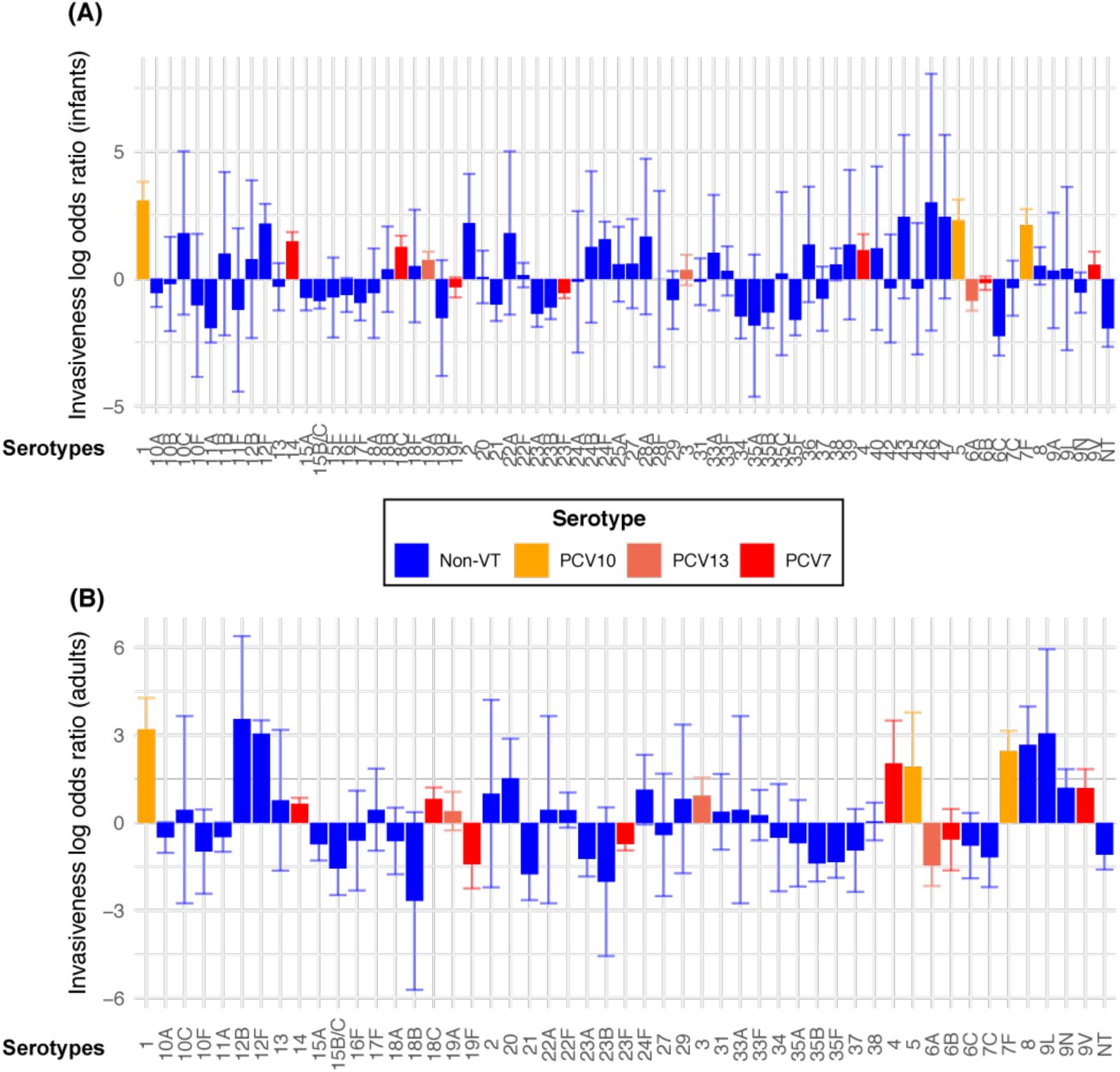
Variation in invasiveness between serotypes. These barcharts show the logarithmic invasiveness odds ratios calculated from the meta-analysis of IPD and carriage isolates (Table S1, S2). The 95% confidence intervals associated with these estimates are shown by the associated error bars. Results are coloured according to the currently-available vaccines in which the serotype is found, if any. **(A)** Invasiveness in infants (those under five) relative to carriage in infants. **(B)** Invasiveness in adults (those over five) relative to carriage in infants. Fewer serotypes are present in this panel, as there were fewer datasets available to estimate these values (Tables S1, S2).

**Fig. S6.**
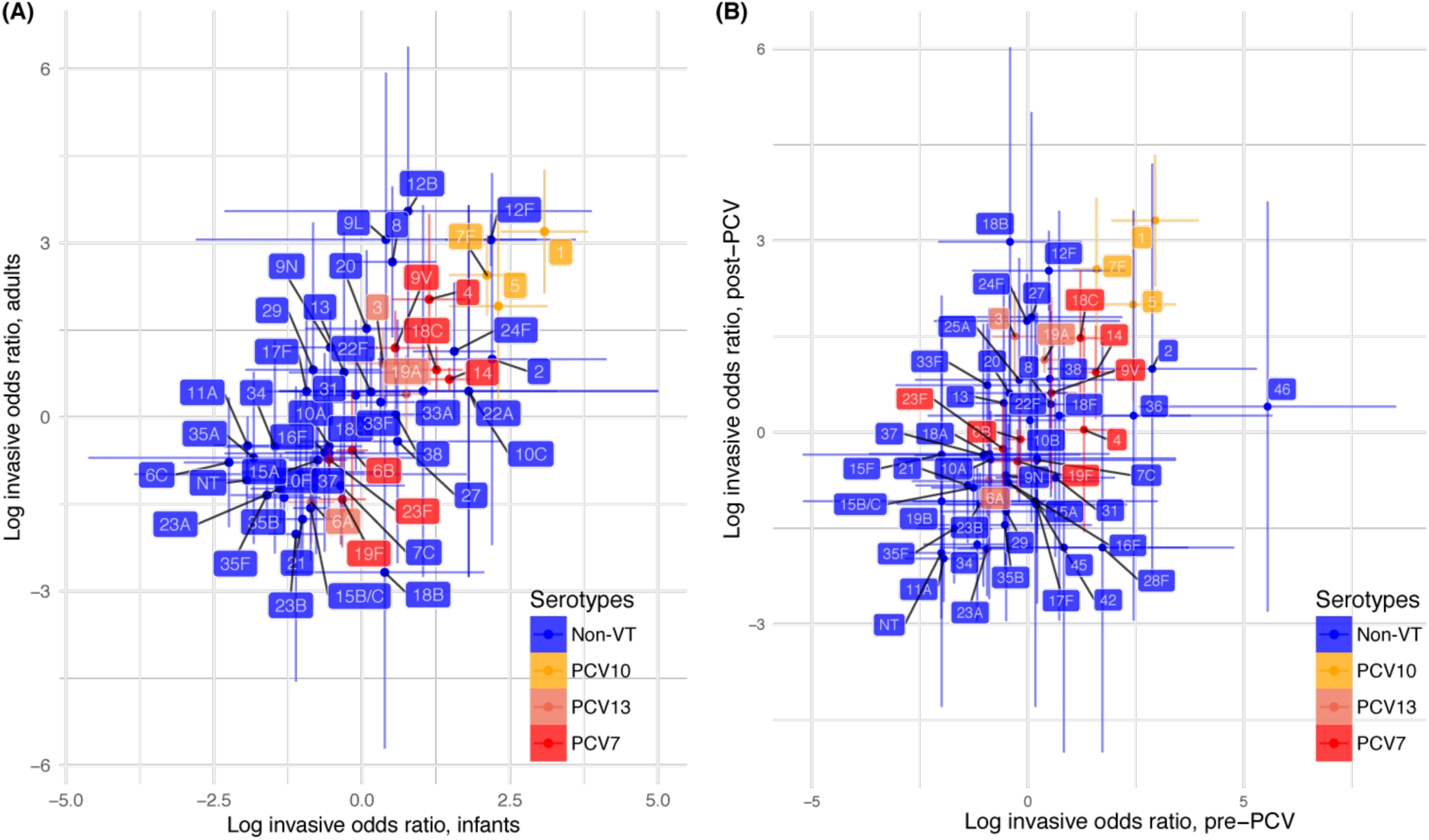
Relationships between serotype invasiveness estimates. **(A)** Invasiveness in infants and adults. This shows the same data as in Fig. 1A, but with all serotypes labelled. **(B)** Invasiveness measures pre- and post-PCV introduction. This plot compares the estimates of the logarithmic odds ratio of invasiveness from the meta-analysis, split by pre- or post-PCV introduction. Considerable variation is evident between the two periods, but the vaccine serotypes do not show particularly high levels of difference. This suggests PCVs do not have a substantial effect on serotype invasiveness. Therefore simulations are justified in associating the same invasiveness with a serotype, regardless of whether it is in the selected PCV formulation or not.

**Fig. S7.**
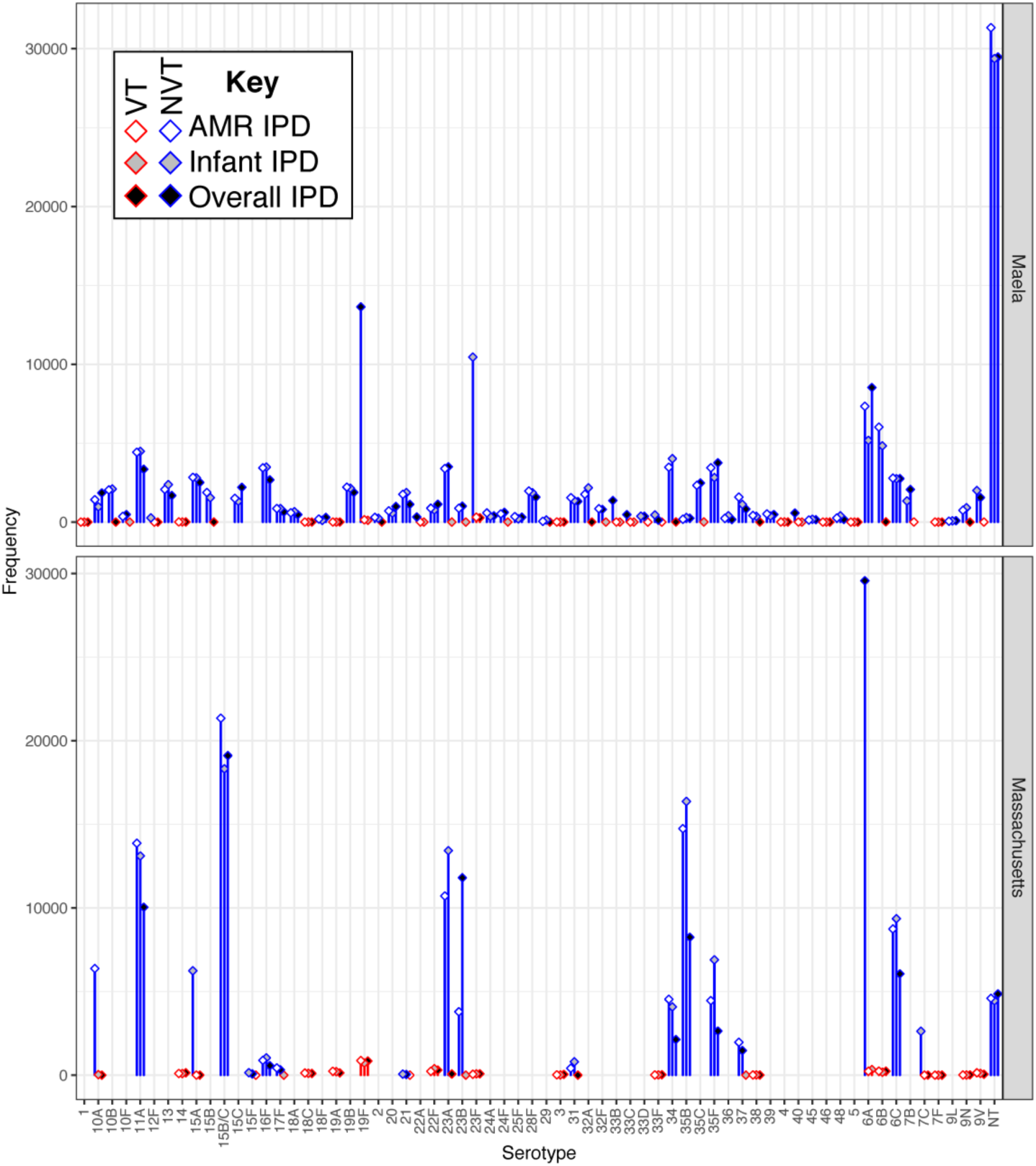
Differences in serotype prevalences 10 years after vaccine introduction between the best-performing 20-valent strategies optimised under different criteria in the two locations. Bars are coloured according to whether they represent the frequency of a vaccine serotype in the corresponding formulation. In Massachusetts, serotypes 6C, 11A, 15B/C and 35B are typically prevalent regardless of the optimisation criterion, owing to their low infant invasiveness. Serotypes 15A and 23A are higher when minimising infant IPD, whereas serotypes 6A and 23B are higher when minimising overall IPD, in accordance with their age-specific invasiveness (Fig. S6). Minimising AMR IPD results in higher prevalence of serotype 10A, which is pansusceptible in Massachusetts. In Maela, all optimal formulations result in serotypes 6A, 6C, 11A, 15F, 19B, as well as non-typeables, remaining at relatively high frequencies in the post-vaccine population. Serotypes 19F and 23F are common when optimising for overall and infant IPD, respectively; both are suppressed when optimising for AMR IPD, owing to their antibiotic resistance profiles. These are partially replaced by serotypes 6A and 6B, which have a weaker association with resistance.

**Fig. S8.**
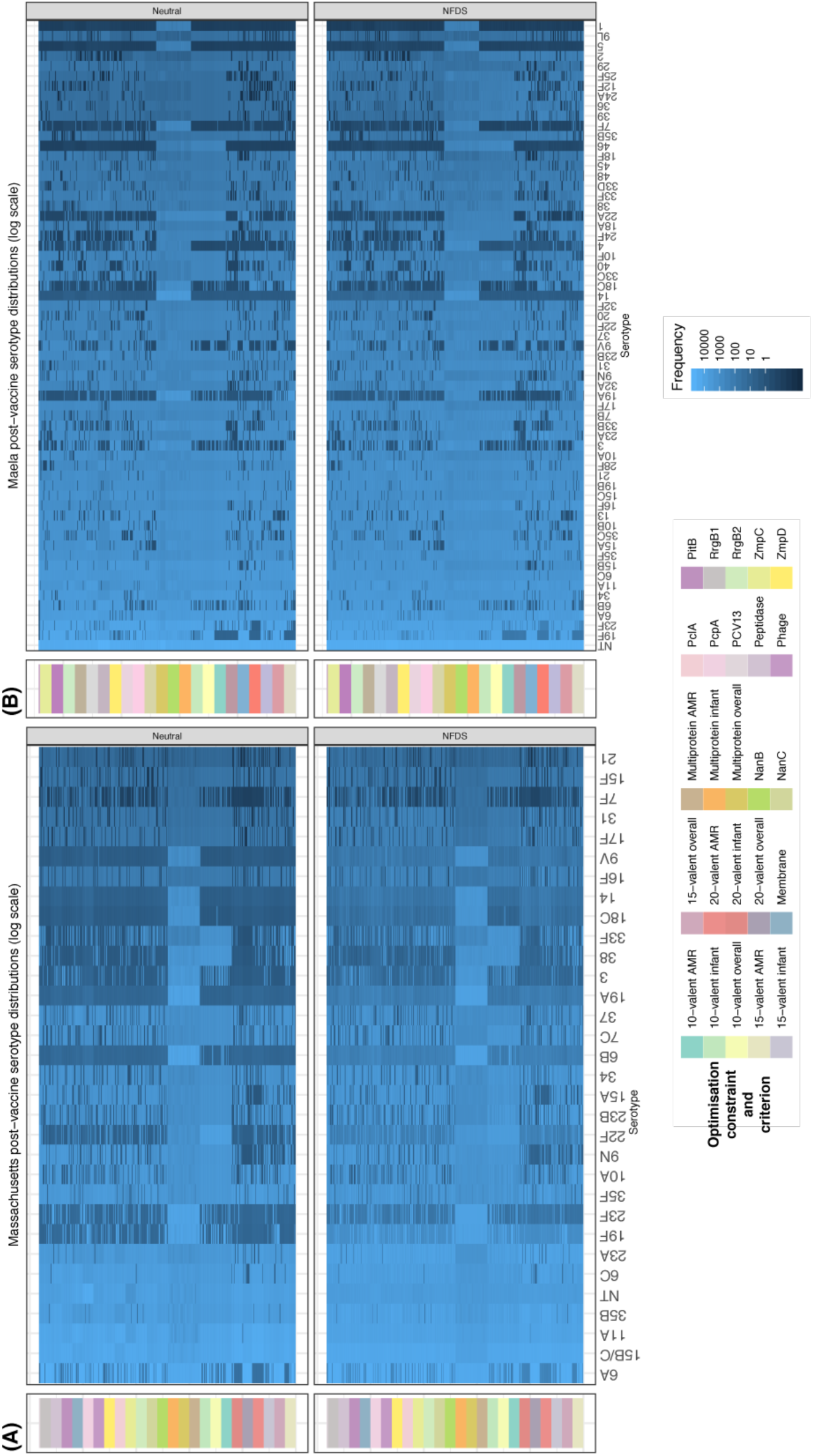
Serotype composition of the post-vaccination populations for all optimised infant-administered vaccination strategies, as indicated by the column on the left. The heatmaps show the simulated frequency of each serotype after 10 years of either multi-locus NFDS, or neutral, evolution on a logarithmic scale for **(A)** Massachusetts and **(B)** Maela.

**Fig. S9.**
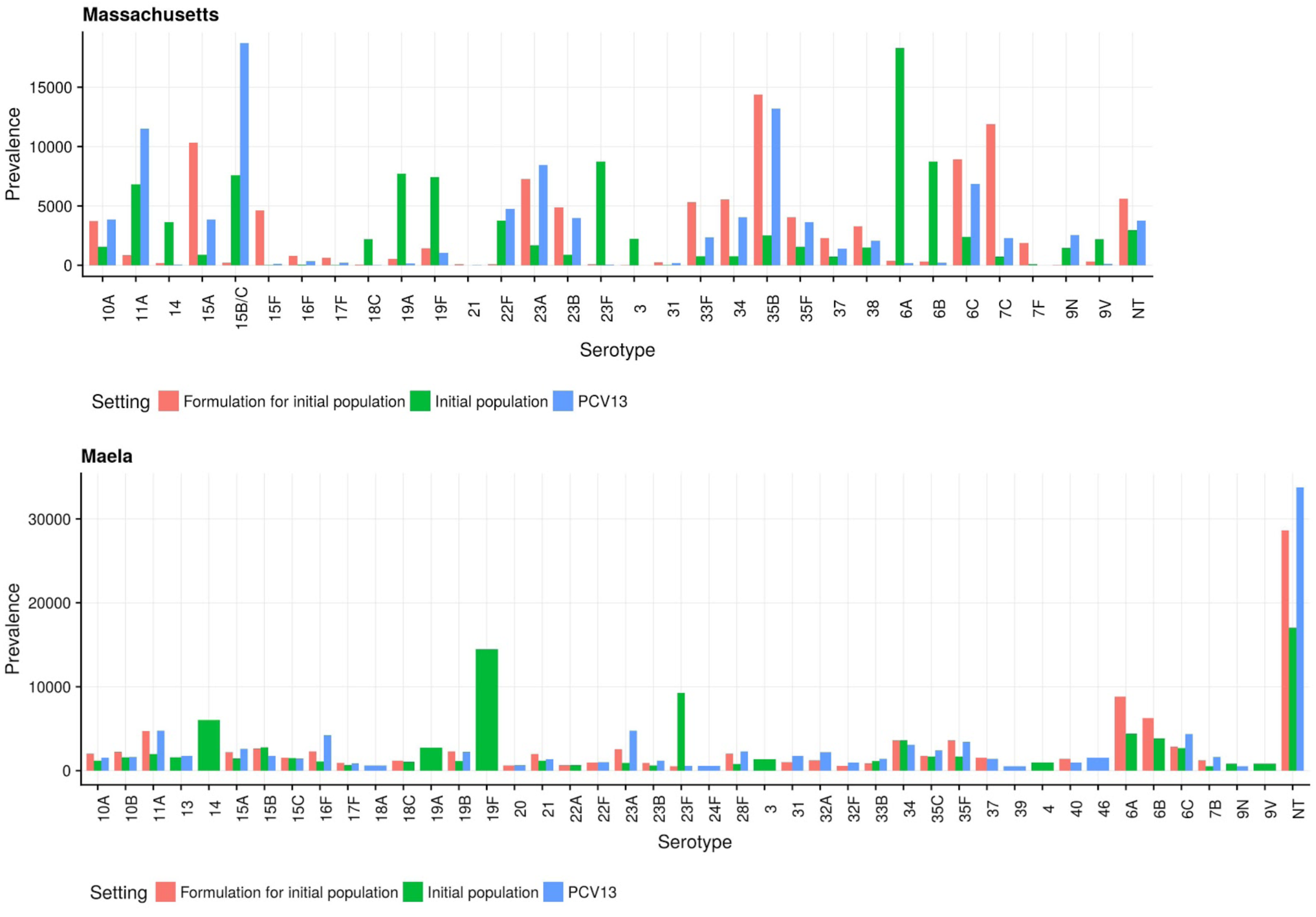
Comparison between the initial modelled populations, formulations containing the serotypes contributing most to infant IPD in the initial population (labelled ‘formulation for initial population’) and the predicted response to the PCV13 vaccine in the two populations. Top: Massachusetts. The model predicts that the ‘formulation for initial population’ strategy would result in an infant IPD burden of 0.64, compared to PCV13’s score of 0.42, and our optimal 15-valent strategy’s score of 0.37. In Maela (bottom) the ‘formulation for initial population’ strategy has an infant IPD burden of 0.59, PCV13 of 0.88, and our optimal 15-valent formulation 0.50. Therefore, in both datasets, our optimised strategies are predicted to out-perform formulations designed based on identifying the serotypes most commonly causing IPD prior to vaccine introduction. Only serotypes with a fraction higher than 0.5% of the simulated population are shown in the Maela population.

**Fig. S10.**
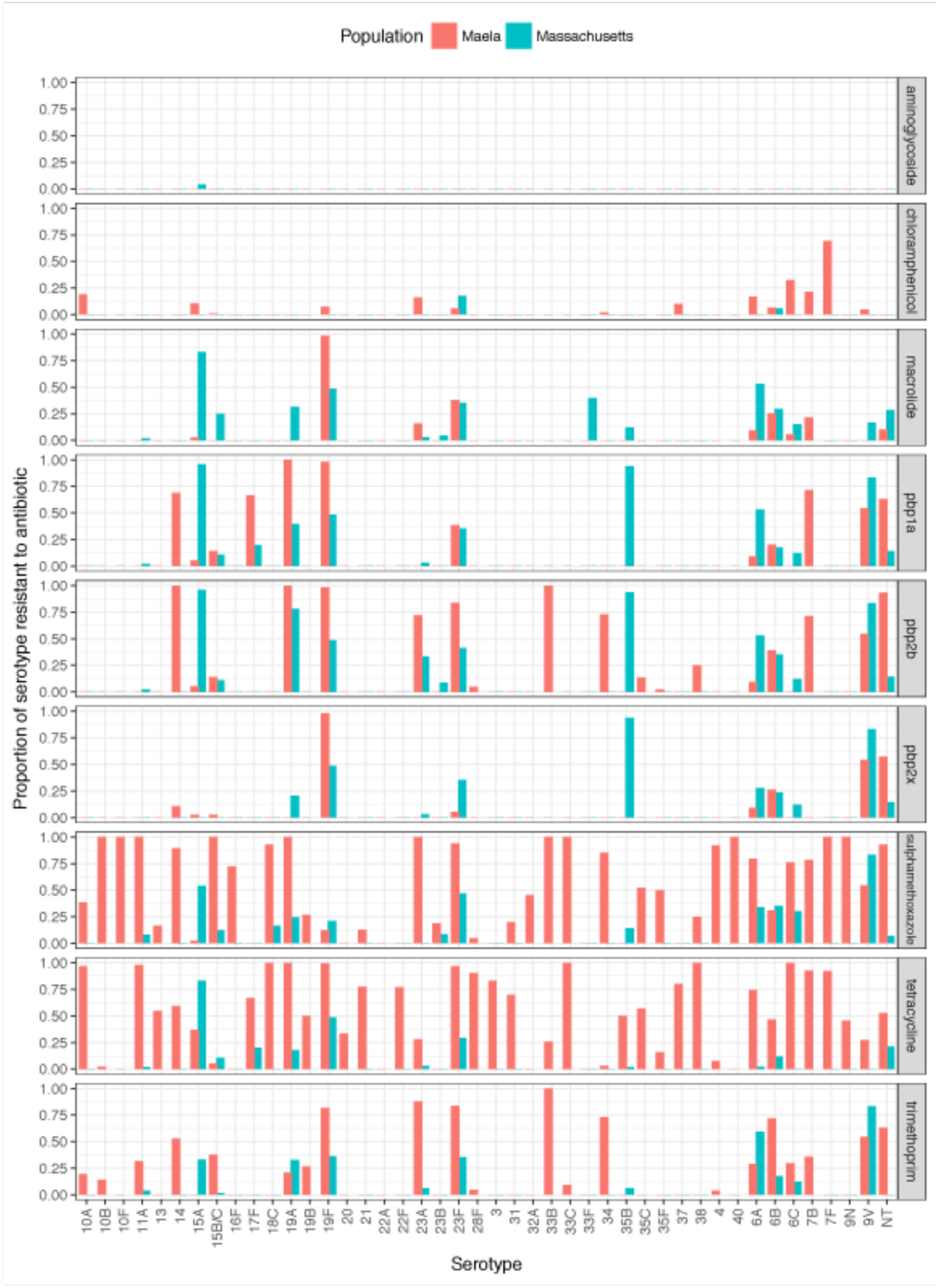
Frequency of resistance loci within each serotype across the Massachusetts and Maela populations.

**Fig. S11.**
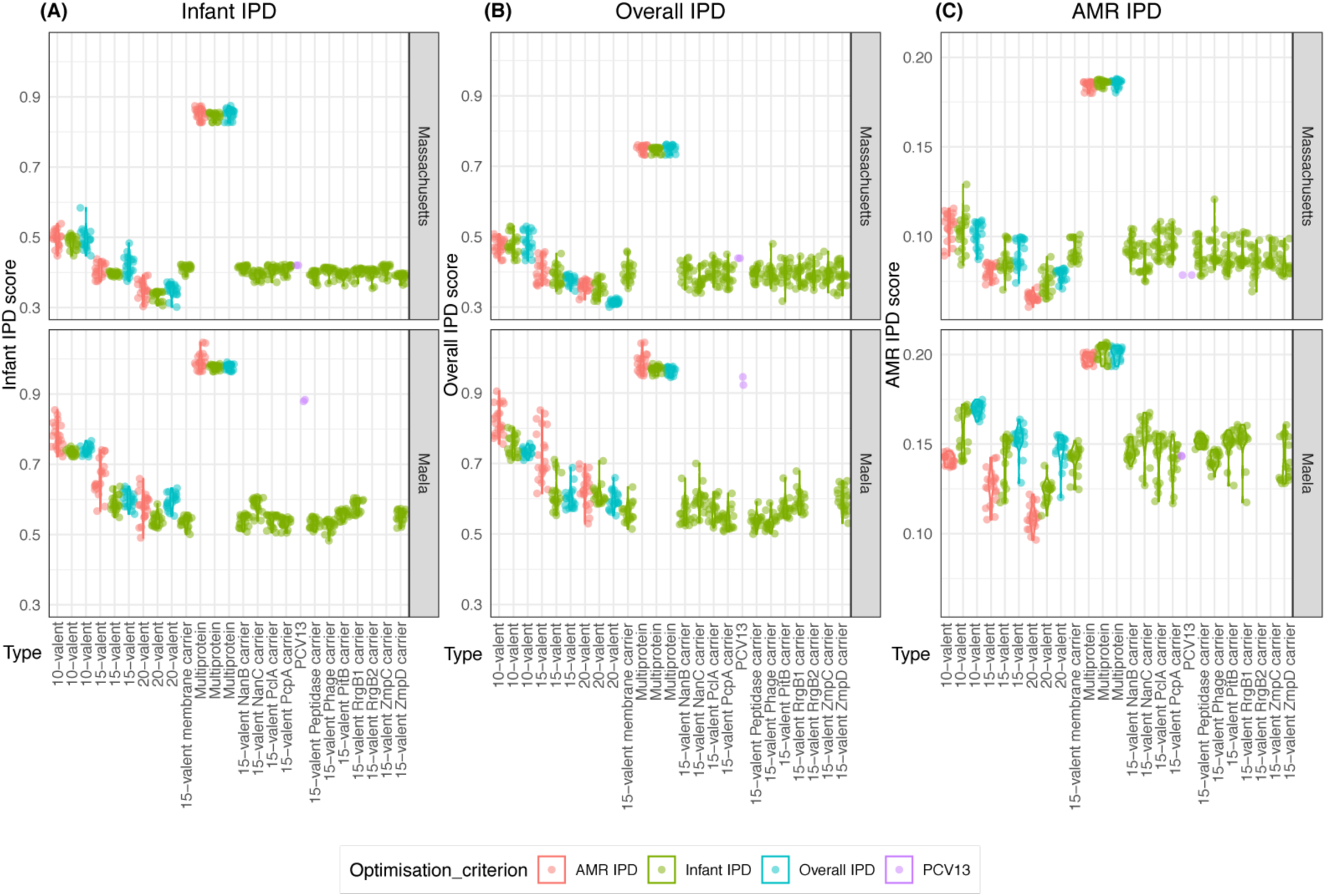
Performance of vaccination strategies judged by different criteria: **(A)** minimising infant IPD; **(B)** minimising overall IPD; **(C)** minimising AMR infant IPD. For the Maela population, no optimisation was performed for two proteins (RrgB2 and ZmpC) that were below the threshold frequency of 0.05 in the starting population (Fig. S12), and therefore not included in the multi-locus NFDS simulations.

**Fig. S12.**
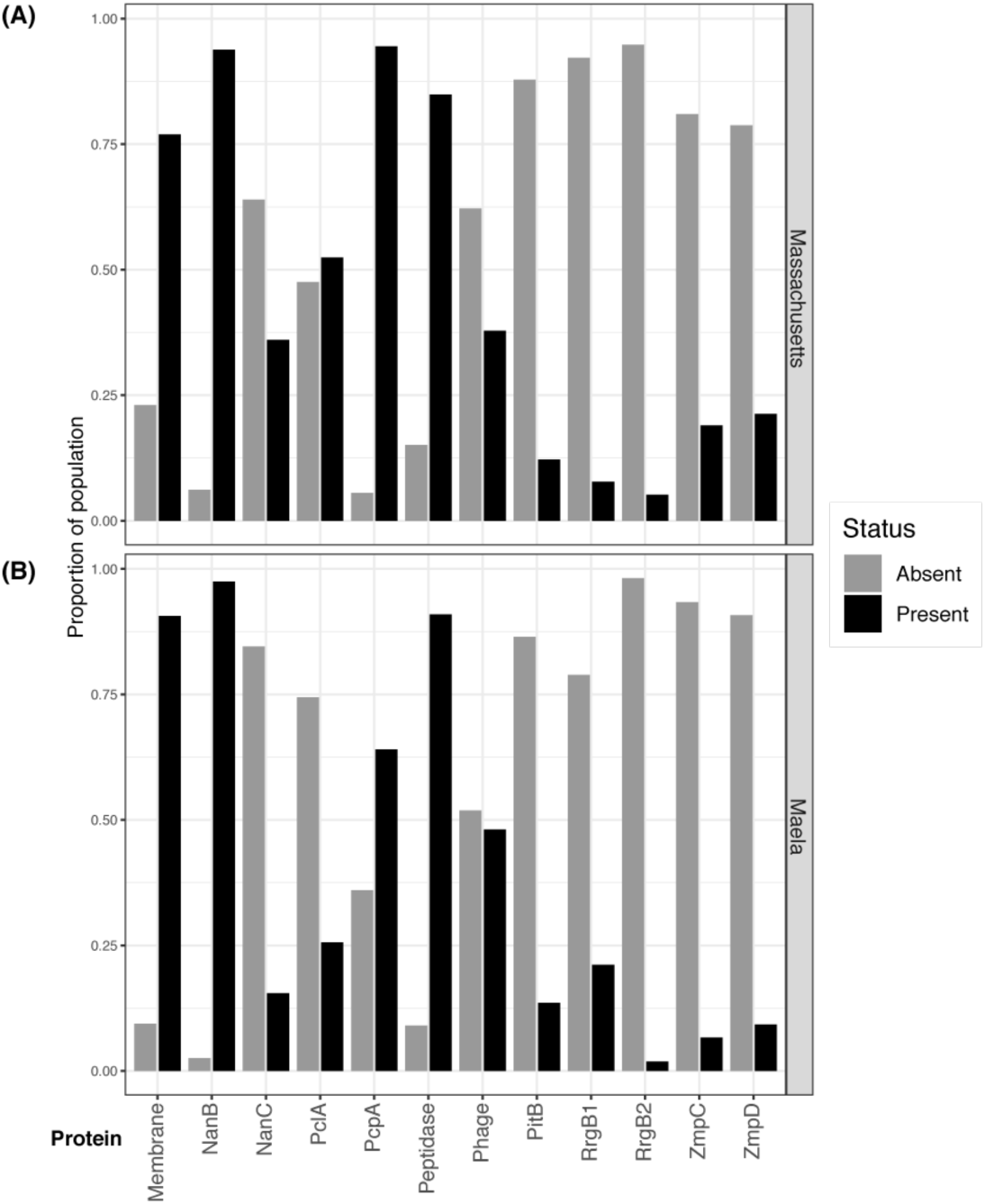
Frequencies of the variable protein antigens in the two pneumococcal populations. These show isolates both exhibiting, and lacking, the antigen co-circulate in the same population. Therefore vaccine-induced immunity against these antigens might facilitate replacement by antigen-negative conspecific competitors.

**Fig. S13.**
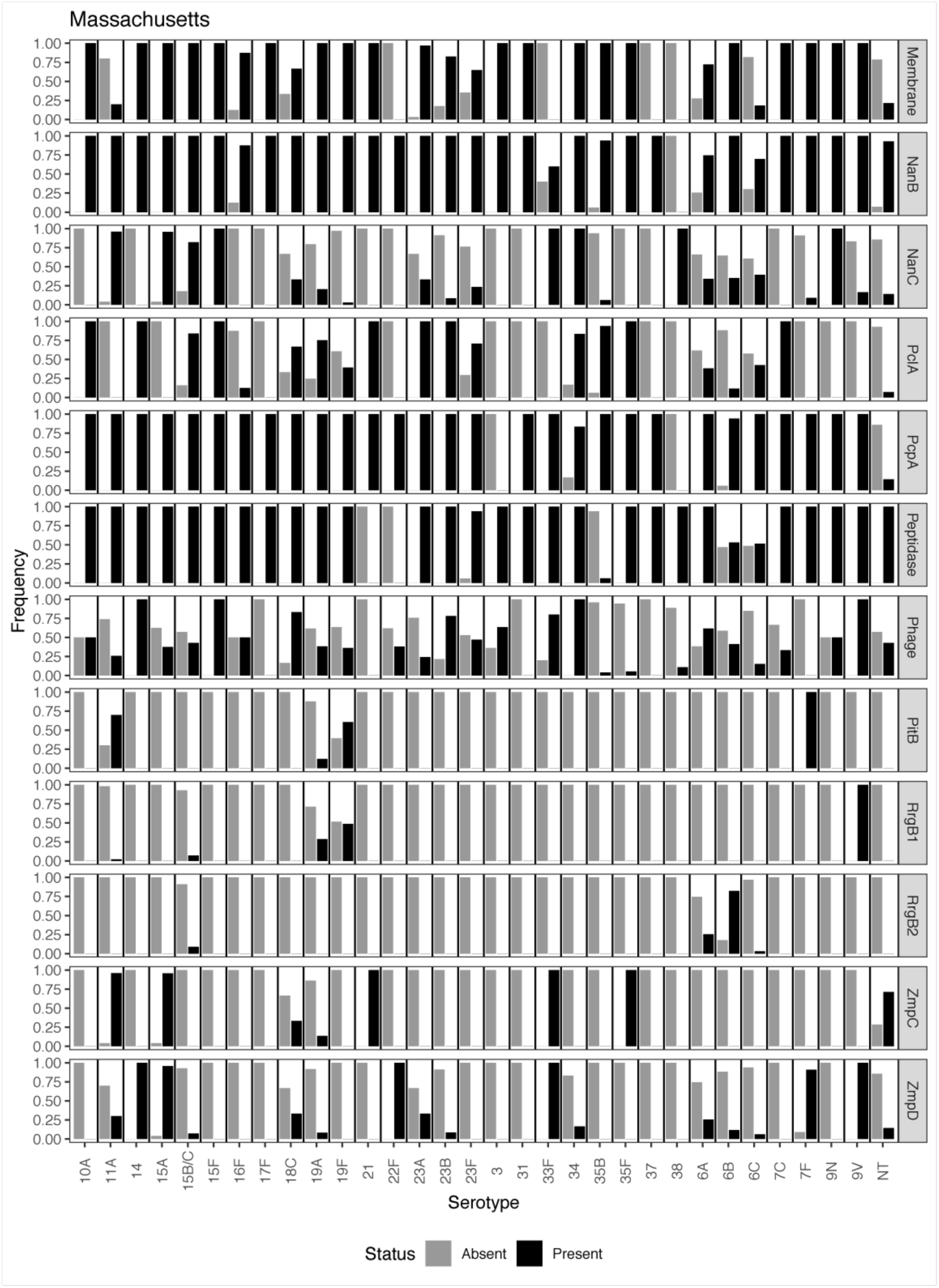
Distribution of protein antigens relative to serotypes in the Massachusetts pneumococcal population.

**Fig. S14.**
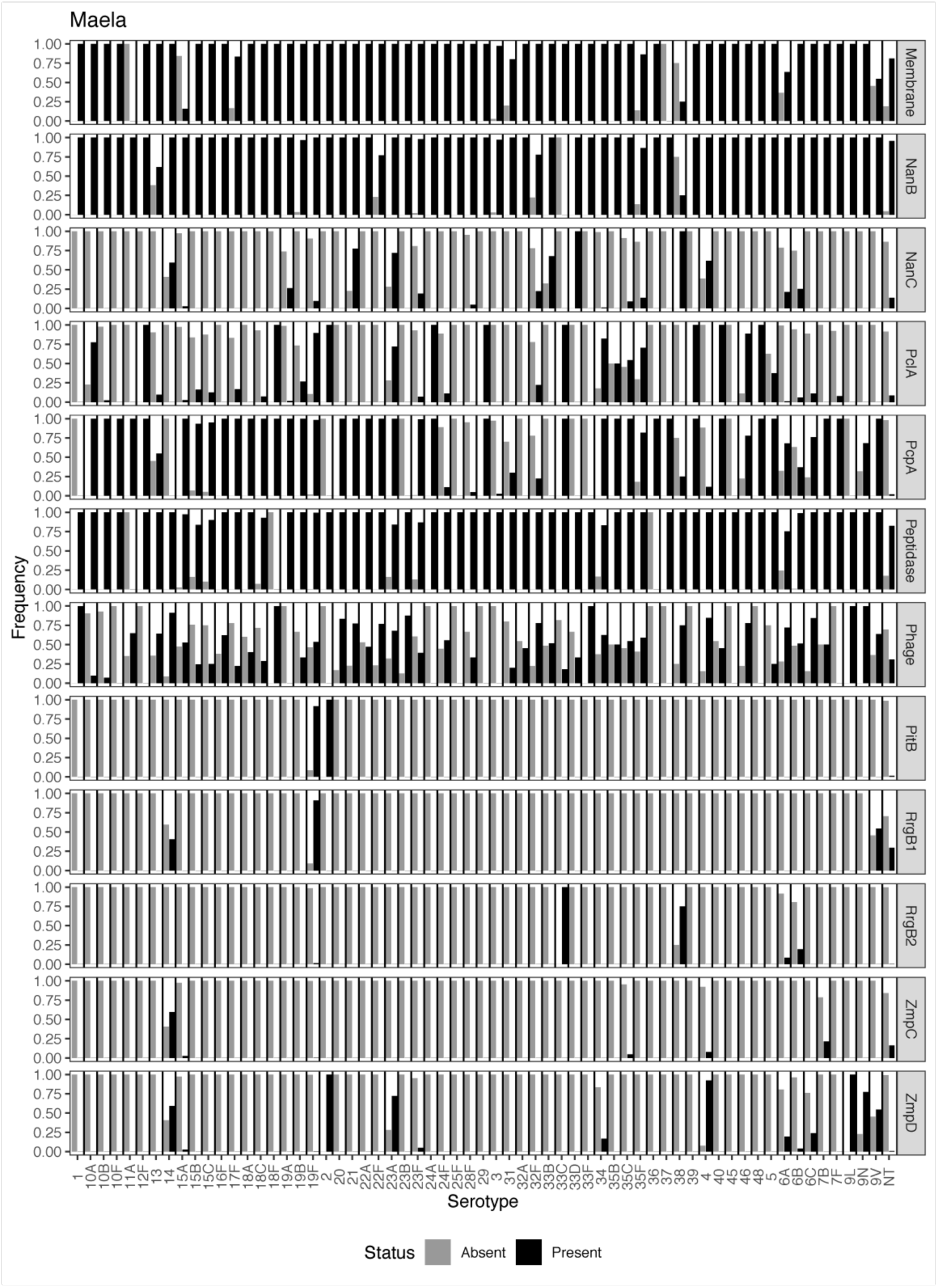
Distribution of protein antigens relative to serotypes in the Maela pneumococcal population.

**Fig. S15.**
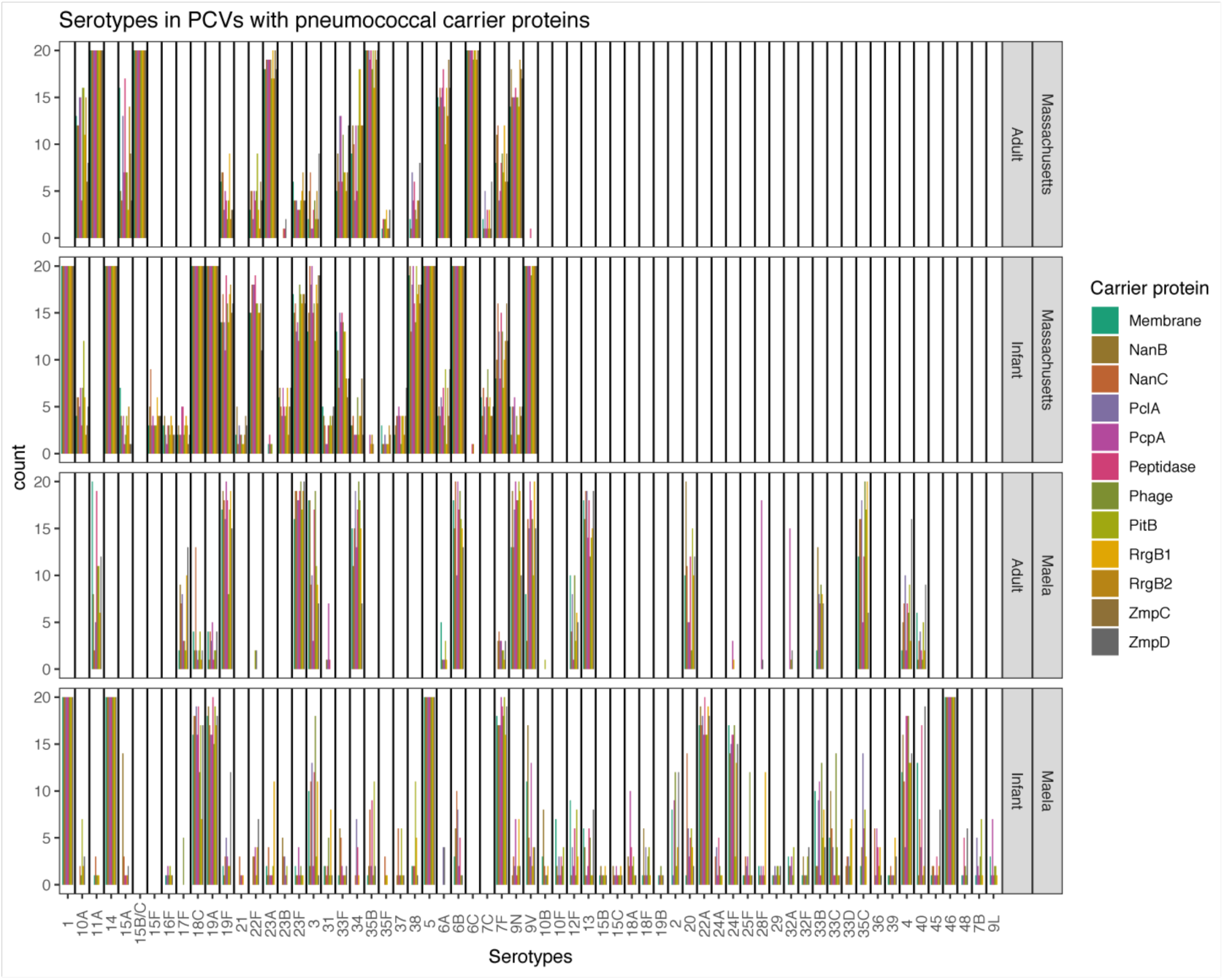
Distribution of capsular antigens between vaccine formulations with pneumococcal carrier proteins. Each bar chart shows the frequency of each capsule type in the 20 optimised formulations for each pneumococcal carrier protein. Panels are split by vaccinee demographic and location.

**Fig. S16.**
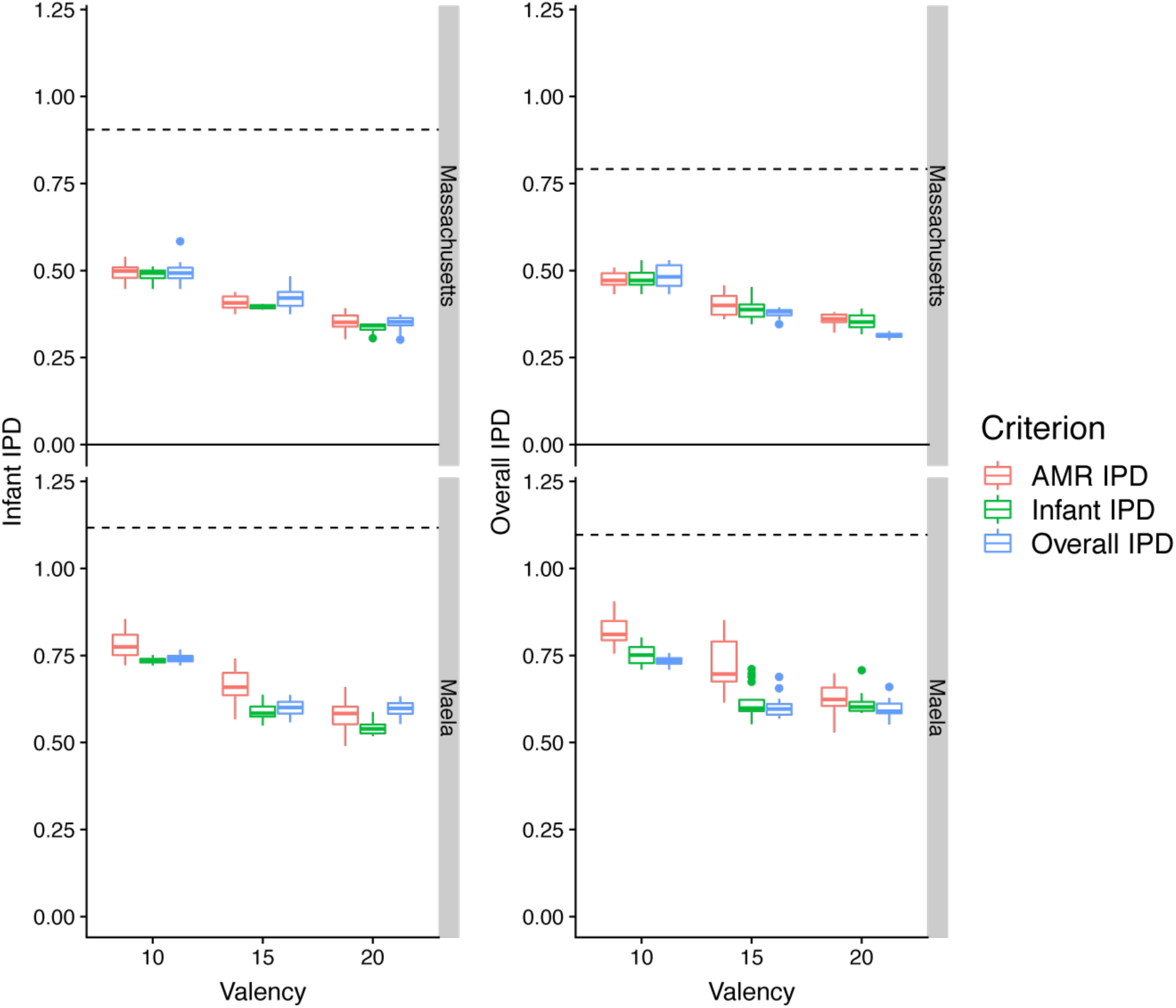
Diminishing returns of expanding PCV valency. Each plot shows the 10 year post-vaccine IPD burden estimated for PCVs of different valencies (including serotypes 1, 5 and 14 in the counts). The box colours show the criterion for which the PCV was optimised. The horizontal dashed line shows the pre-vaccination IPD burden in the relevant population. The reduction in IPD caused by the 20-valent PCVs is not double that achieved by the 10-valent PCVs, despite the latter being constrained to only the serotypes present in PCV13.

**Fig. S17.**
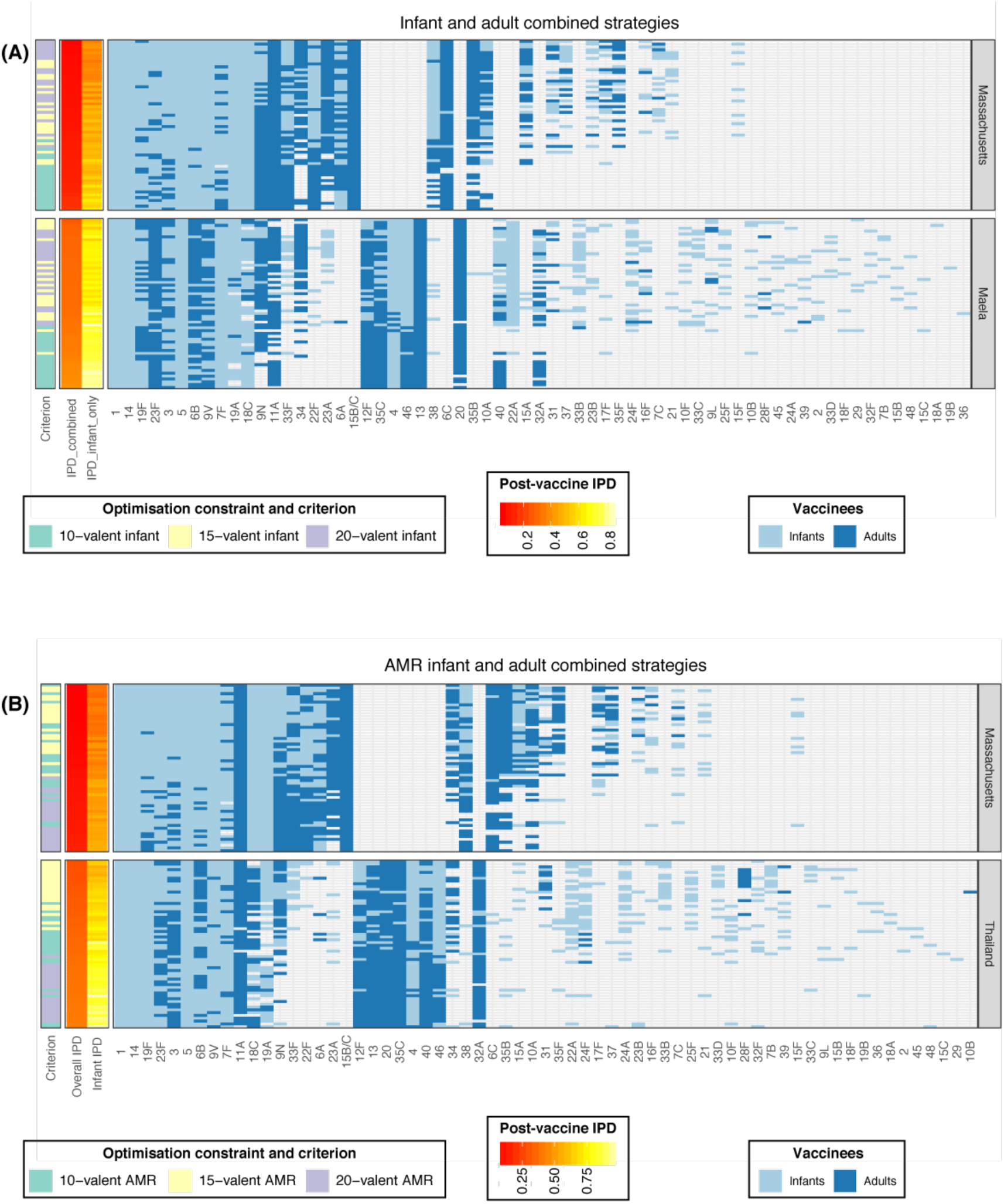
Combined vaccination strategies for minimising IPD. For each infant-administered PCV design, a complementary adult vaccine was identified to target the 10 serotypes predicted to cause the highest levels of post-PCV IPD in adults. On each row, the light blue cells define the infant formulation, and the dark blue cells define the adult formulation. Rows are ordered by the overall IPD burden estimated following the implementation of the combined vaccination strategy. **(A)** Combined strategies in which the infant-administered vaccine minimised infant IPD (corresponding to the vaccines in Fig. 1D,E), and the adult-administered vaccine minimised residual adult IPD. **(B)** Combined strategies in which the infant vaccine minimises overall AMR IPD (corresponding to the vaccines in Fig. 3D,E), and the adult vaccine minimises residual adult IPD.

**Fig. S18.**
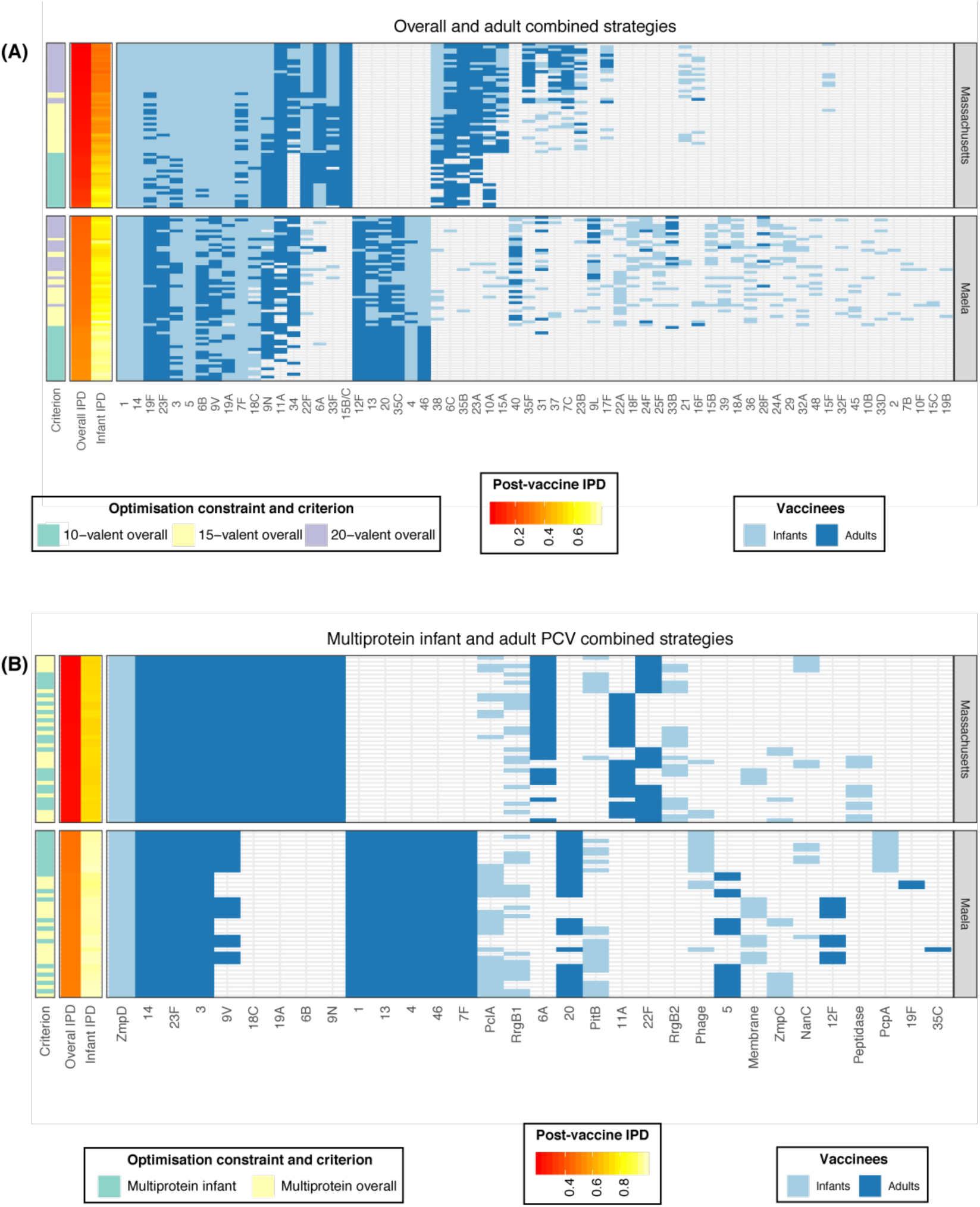
Further combined vaccination strategies for minimising IPD, displayed as described in Fig. S17. **(A)** Combined strategies in which the infant vaccine minimises overall infant and adult IPD, and the adult vaccine minimises residual adult IPD. **(B)** Combined vaccination strategies in which a PCV for use in adults is designed to be complementary to the multiprotein infant vaccine. Complementarity is exemplified by the “Membrane” protein-based formulations. In Maela, highly invasive serotype 12F isolates do not express this protein (Fig. S14), and hence this serotype is present in the adult vaccines complementary to “Membrane” protein-based infant vaccines.

**Fig. S19.**
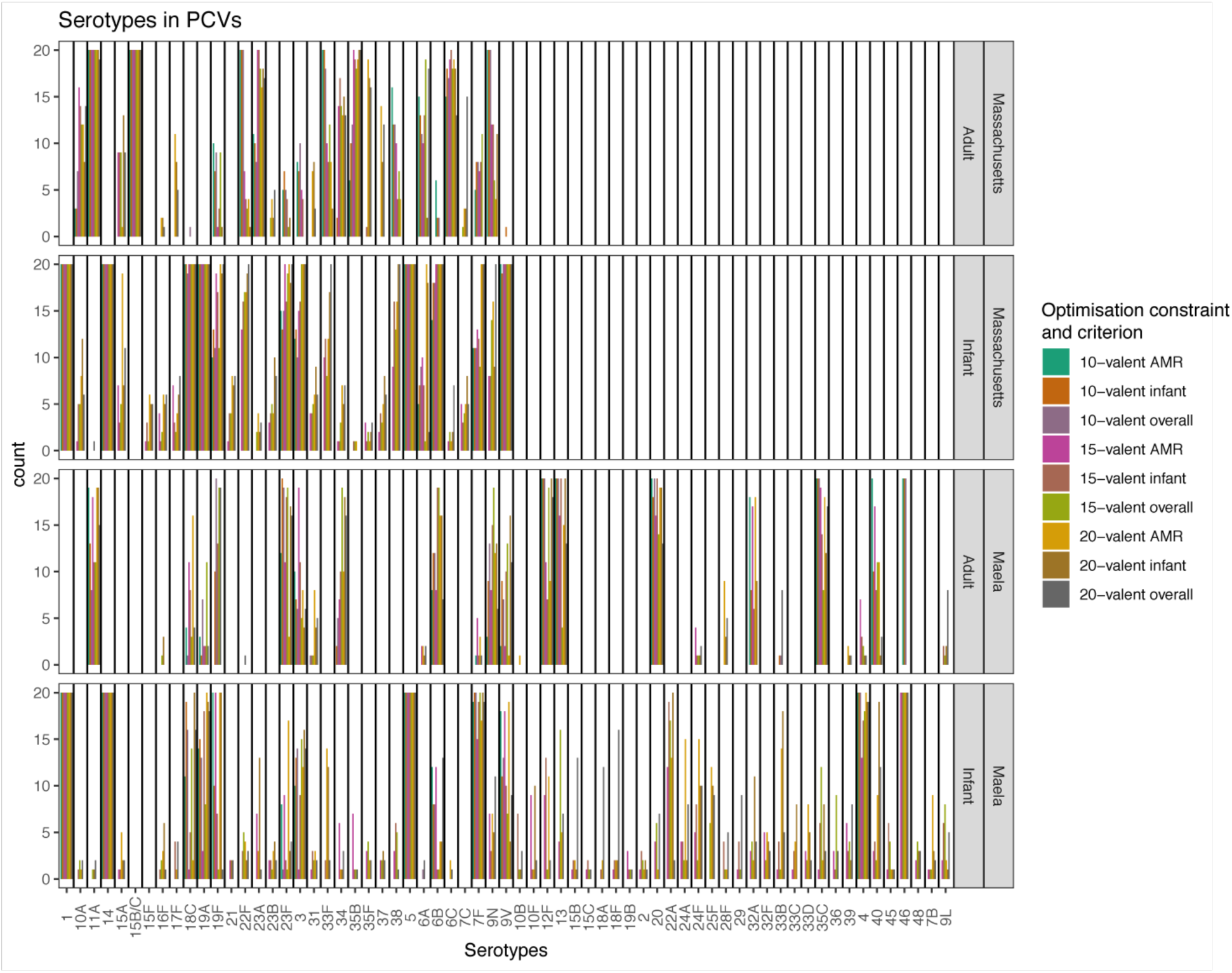
Distribution of capsular antigens between vaccine formulations. Bar charts show the frequency of each capsule type in the 20 analysed formulations for each combination of criterion and constraint under which optimisation was performed, as represented by the bar colour. Panels split the formulations by population (Massachusetts or Maela) and vaccines administered to infants, and the complementary adult vaccines (CAVs). Data are displayed as in Fig. S15.

**Fig. S20.**
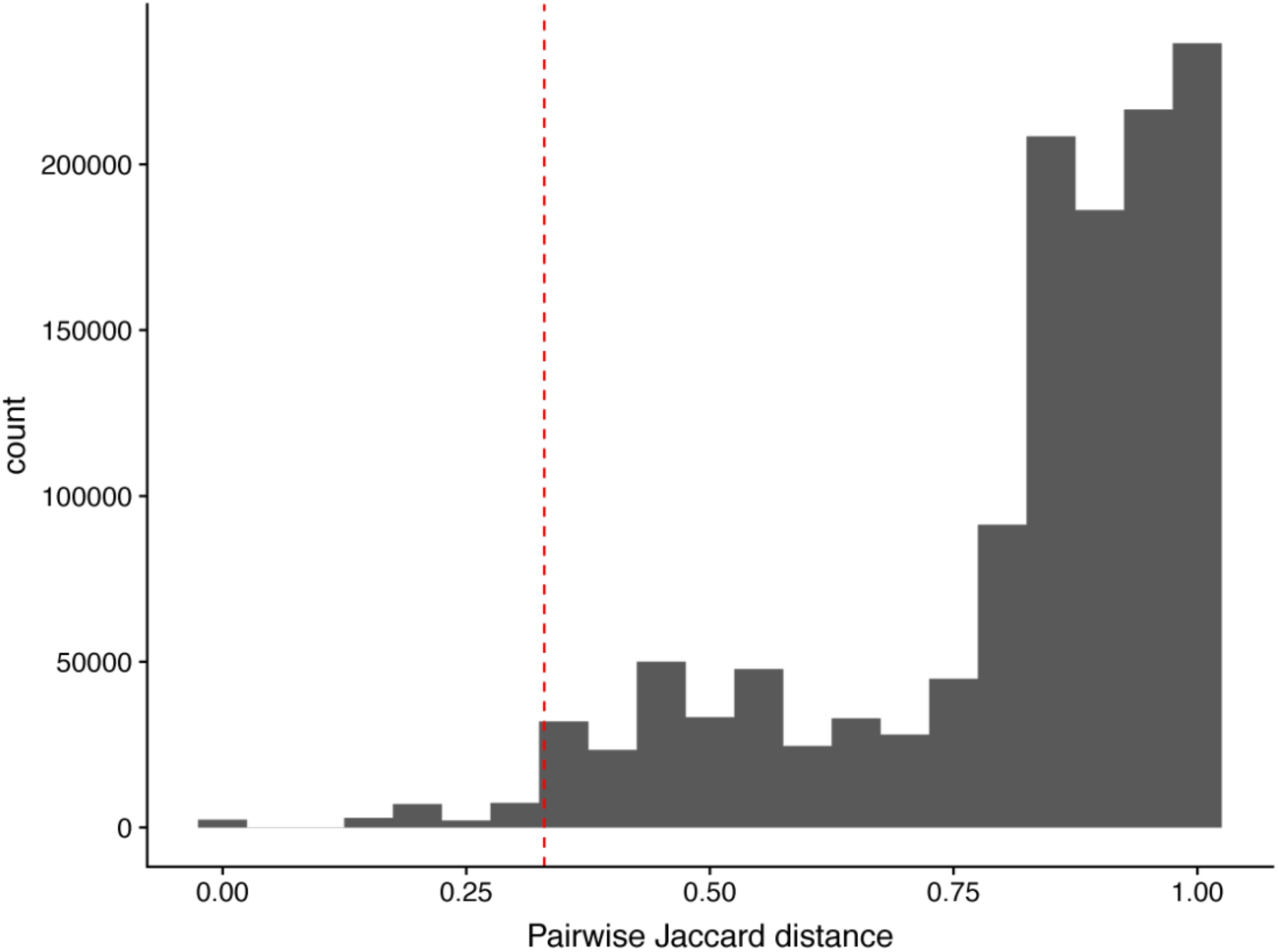
Distribution of pairwise Jaccard distances between vaccine formulations. The vertical red dashed line shows the threshold similarity (0.33) used to define edges in the network displayed in Fig. 5C.

**Fig. S21.**
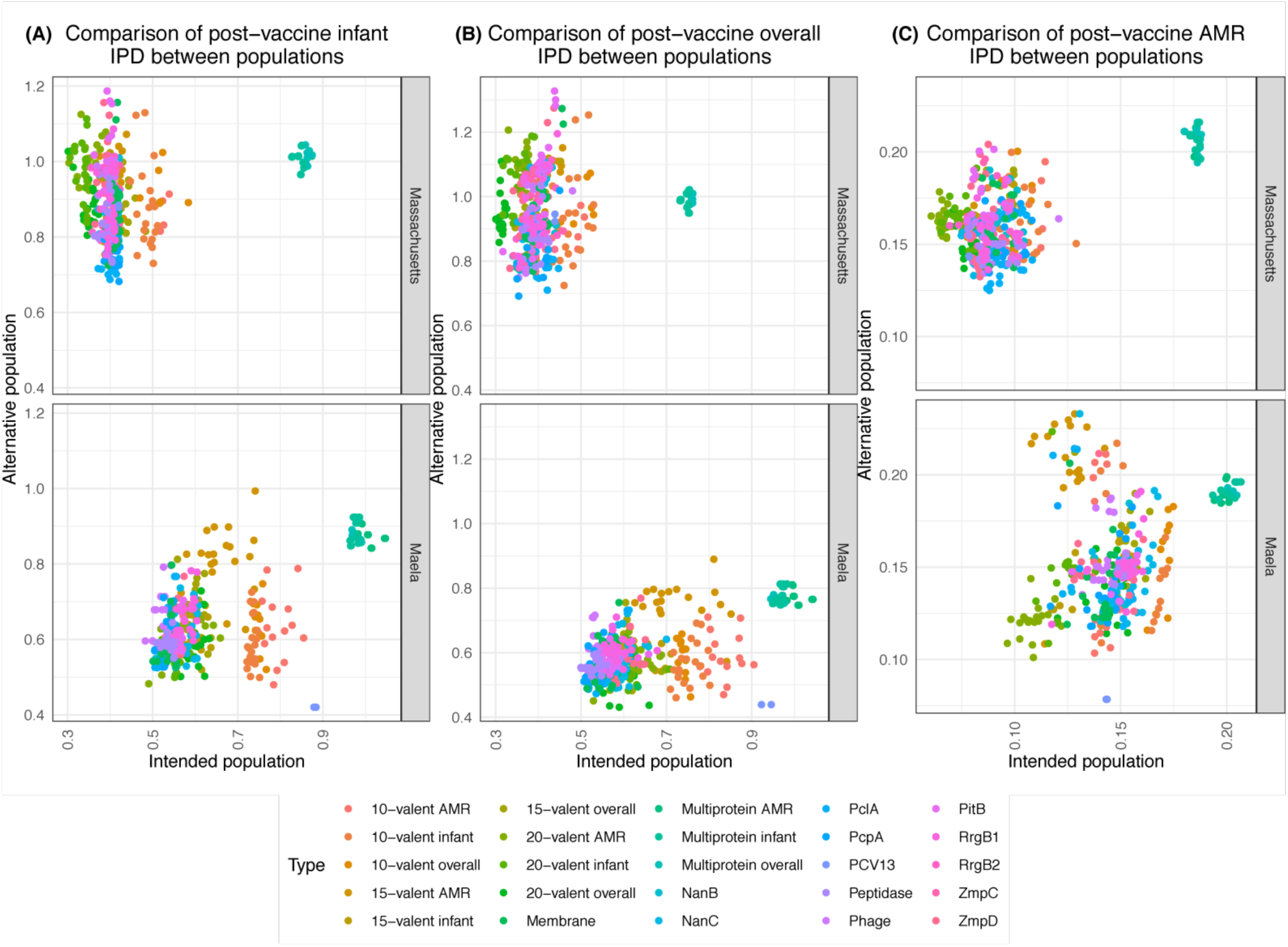
Performance of vaccine strategies in the alternative population to that for which they were designed. Panels are labelled to indicate the population for which the formulation was designed. Simulations of each strategy were run in the alternative population, and the performance assessed by different criteria: **(A)** minimising infant IPD; **(B)** minimising overall IPD; **(C)** minimising AMR infant IPD. Notably, those vaccines designed to reduce infant and overall IPD in Massachusetts are predicted to perform very poorly in Maela.

**Fig. S22.**
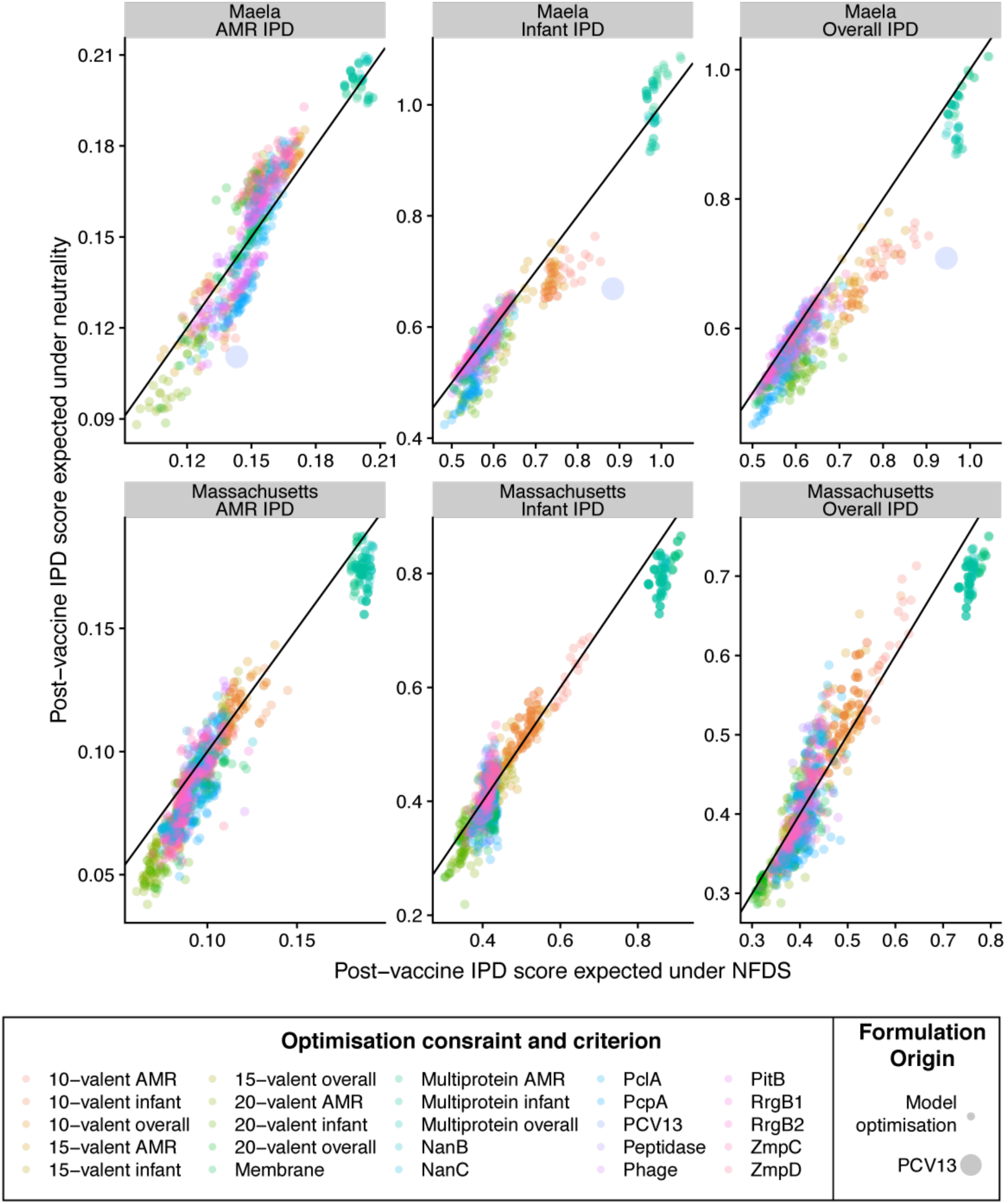
Comparing the simulated effectiveness of vaccine formulations in the original multi-locus NFDS model and an otherwise equivalent neutral model. Each plot shows the expected post-vaccine IPD burden measure expected under NFDS and neutral evolution; points are coloured by optimisation constraint and criterion, and the line of identity is marked. The results correlate strongly, with each optimisation criterion generally predicted to be slightly lower in the neutral model. Vaccine compositions that we predict to perform better than PCV13 tend also to do so in the neutral model. This indicates the formulations we have identified perform well despite the predicted effects of NFDS, rather than because of them.

**Fig. S23.**
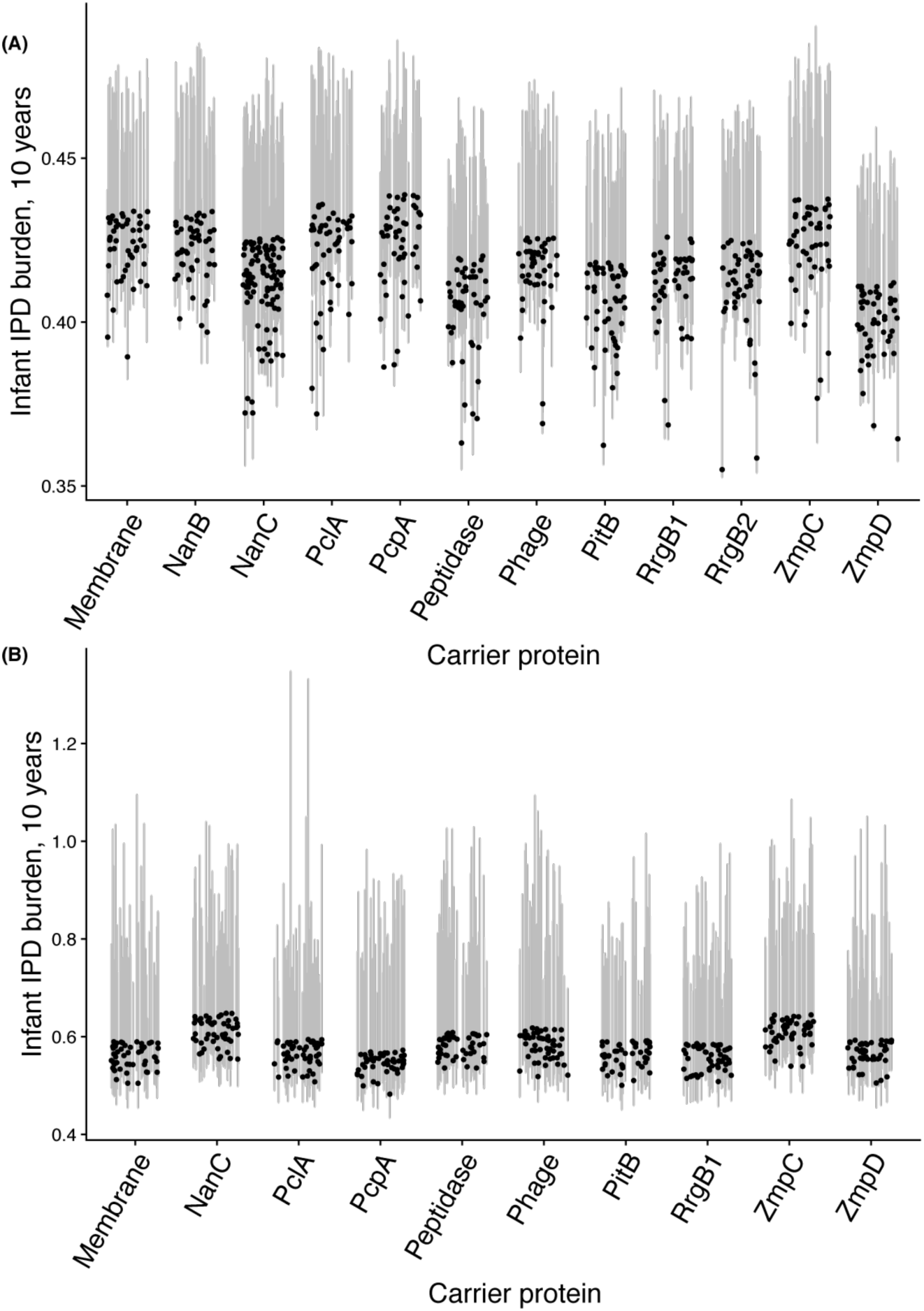
Variation in estimated IPD burden with resampling of serotypes’ invasiveness. The infant IPD burden was calculated for the 15-valent PCVs containing a pneumococcal carrier protein in **(A)** Massachusetts and **(B)** Maela. Proteins including the Peptidase, PitB and ZmpD proteins as carriers consistently achieved a lower point estimate of infant invasiveness than PCV13. Grey lines show inter-quartile ranges; these are positively skewed, due to the Gaussian distribution assumption on the invasiveness logarithmic odds ratios combined with the use of non-logarithmic odds ratios in the optimisation criteria. The uncertainty is greatest for serotypes rarely included in epidemiological studies, with the consequence that the Maela estimates are associated with much greater uncertainty than the Massachusetts estimates.

## Supplementary Tables

**Table S1. (separate file)**

Epidemiological studies included in the meta-analysis of age-specific serotype invasiveness.

**Table S2. (separate file)**

Epidemiological data for the meta-analysis of age-specific serotype invasiveness.

**Table S3.** Characteristics of the intermediate-frequency *S. pneumoniae* protein antigens. Each protein antigen is listed by its descriptor and the corresponding cluster of orthologous genes in Corander *et al* ^12^ and Croucher *et al* ^10^; the sequences of all proteins in the latter are available from http://datadryad.org/resource/doi:10.5061/dryad.t55gq. Most of these proteins were identified using a panproteome array, but others were previously discovered by more targeted approaches.

| Descriptor | Cluster of orthologous genes in Croucher *et al* | Cluster of orthologous genes in Corander *et al* | Function | Evidence for immunogenicity |
| --- | --- | --- | --- | --- |
| NanB | CLS01445 | CLS00257 | neuraminidase B | 36 |
| ZmpD | CLS02608 | CLS00476 | zinc metalloprotease D variant | 36 |
| PclA | CLS03178 | CLS00440 | pneumococcal collagen-like protein A variant | 36 |
| RrgB1 | CLS02942 | CLS02709 | type I pilus rrgB (clade 1) structural protein | 63 |
| PitB | CLS02871 | CLS01706 | type 2 pilus structural protein PitB | 36 |
| RrgB2 | CLS02796 | CLS03842 | type I pilus rrgB (clade 2) structural protein | 63 |
| Phage | CLS01887 | CLS00695 | Prophage protein | 36 |
| Membrane | CLS00011 | CLS01683 | Membrane protein of unknown function | 36 |
| ZmpC | CLS01991 | CLS04319 | zinc metalloprotease C | 36 |
| NanC | CLS01160 | CLS03670 | neuraminidase C | 36 |
| Peptidase | CLS01541 | CLS01895 | M50 peptidase family protein | 36 |
| PcpA | CLS01852 | CLS01587 | choline binding protein PcpA | 36 |

**Table S4.** Common features of optimized vaccine formulations. For each of the demographics (infant and adult) and regions (Massachusetts and Maela), these descriptions define the common serotypes included in the optimised formulations identified when minimizing the burden of infant, overall or AMR IPD. These were identified through logic regression against a random set of formulations, followed by manual curation to generate more intuitive descriptions.

| Vaccinee demographic and region | Common features of formulations |
| --- | --- |
| Massachusetts infants | Contains a core of 1, 5, 18C, 14, and 19A; plus at least one of 6B or 9V; plus at least three of 19F, 6A, 23F, 3, 38, 7F, 33F, 22F |
| Massachusetts adults | Contains a core of 11A, 15B/C; plus one of 23A, 6C, 9N or 10A; plus one of 35B, 6A, 33F |
| Maela infants | Contains a core of 1, 14, 46 and 5; plus four of 24F, 22A, 40, 4, 10F, 7F, 19A, 18C, 9L, 19F, 35C, 3, 33C, 9V, 23B, 15A, 15B, 36, 32A, 45, 15A, 16F<br>OR<br>Contains a core of 1, 14, 4, 5; plus one of 18C, 19F, 7F, 9V, 19A, 6B, 3 |
| Maela adults | One of 24A, 21, 40, 13, 45; plus four of 23F, 13, 9N, 19F, 35C, 6B, 20, 3, 9V, 34 |

