## Supplementary material for "Designing ecologically-optimised vaccines using population genomics": Table S1 (supplemental)

| **Population** | **Vaccination status** | **Time frame** | **Carriage isolate source** | **Infant disease source** | **Adult disease source** | **References** |
| --- | --- | --- | --- | --- | --- | --- |
| Alabama | Pre-PCV | July 1975 - December 1978 | Unvaccinated sick and healthy children <18 years old from Alabama | Unvaccinated children with IPD <18 years old from Alabama | Unvaccinated adults hospitalised with pneumonia or IPD in Alabama | (Gray et al. 1979) |
| Goroka | Pre-PCV | 1981 - 1987 | Unvaccinated children attending clinics near Goroka Town | Unvaccinated children with IPD in Goroka hospital | - | (Smith et al. 1993) |
| Kenya | Pre-PCV | 1992 - 1996 | Unvaccinated sick children in Kilifi <18 years old | Unvaccinated children <12 years old hospitalised with pneumonia or IPD in Kilifi | Unvaccinated adults hospitalised with pneumonia or IPD in Kilifi or Mombasa | (Scott et al. 1998) |
| Iceland | Pre-PCV | 1992-2001 | Unvaccinated children <5 years old | Unvaccinated children <5 years old | - | (Brueggemann et al. 2004) |
| Oxford | Pre-PCV | 1994-2001 | Unvaccinated healthy children <5 years old in Oxford | Unvaccinated children <5 years old with IPD in Oxford | - | (Brueggemann et al. 2003) |
| Finland | Pre-PCV | 1994-1999 | Unvaccinated healthy children <2 years old in Tampere | Unvaccinated children with IPD <2 years old across Finland | - | (Hanage et al. 2005) |
| Ontario | Pre-PCV | 1995 | Unvaccinated healthy children primarily <4 years old in Toronto | Unvaccinated children <18 years ld with IPD in Toronto and surrounding area | - | (Kellner et al. 1998) |
| Atlanta | Pre-PCV | January 1995 - December 1995 | Unvaccinated children with recent URI in Atlanta <5 years old | Unvaccinated children <5 years old with IPD in Atlanta | - | (Sharma et al. 2013) |
| Czech | Pre-PCV | 1996 – 2005 | Unvaccinated healthy children 3 – 5 years old across the Czech Republic | Unvaccinated children <6 years old with IPD across the Czech Republic | - | (Zemlickova et al. 2010) |
| E&W | Pre-PCV | July 1996 – June 2006 | Unvaccinated healthy children <5 years old in Hertfordshire | Unvaccinated children <5 years old across England & Wales | Predominantly unvaccinated individuals >4 years old across England & Wales | (Trotter et al. 2010) |
| Stockholm | Pre-PCV | 1997 | Unvaccinated healthy children <7 years old in Stockholm | - | Unvaccinated adults (and a small number of children) hospitalised with IPD in Stockholm | (Sandgren et al. 2004) |
| Sweden | Pre-PCV | 1997 - 2008 | Unvaccinated children <18 years old from the Stockholm area | Predominantly unvaccinated children with IPD <18 years old from across Sweden | Unvaccinated adults with IPD across Sweden | (Browall et al. 2014) |
| Alaska | Pre-PCV | 1998-2002 | Children <5 years old, likely predominantly unvaccinated | Children <5 years old, likely predominantly unvaccinated | - | (Brueggemann et al. 2004) |
| Greece | Pre-PCV | January 2000 – December 2004 | Unvaccinated children <5 years of age from north-west Greece | Unvaccinated children <18 years of age with IPD from north-west Greece | - | (Levidiotou et al. 2006) |
| Portugal | Pre-PCV | January 2001 - December 2003 | Unvaccinated healthy children <7 years old in Lisbon and Oeiras | - | Unvaccinated individuals, primarily adults, across Protugal | (Sá-Leao et al. 2011) |
| Bogota | Pre-PCV | May 2005 - November 2006 | Healthy unvaccinated children <18 months old in Bogota | IPD in children <2 years old in Bogota | - | (Parra et al. 2013) |
| Caracas | Pre-PCV | December 2006 - January 2008 | Unvaccinated healthy children <6 years old in Caracas | Unvaccinated children with IPD <6 years old in Caracas | - | (Rivera-Olivero et al. 2011) |
| Morocco | Pre-PCV | November 2010 - December 2011 | Unvaccinated healthy children in Rabat | Children hospitalised with severe pneumonia in Rabat < 5 years old | - | (Jroundi et al. 2017) |
| Massachusetts | Post-PCV7 | 2001 – April 2009 | PCV7-vaccinated children visiting physicians <7 years old across Massachusetts | PCV7-vaccinated children <7 years with IPD old across Massachusetts | - | (Yildirim et al. 2010) |
| Navajo | Post-PCV7 | March 2006 – March 2008 | PCV7-vaccinated Navajo or White Mountain Apache native American children <9 years old | Active IPD surveillance of PCV7-vaccinated Navajo or White Mountain Apache native American children <7 years old | Active IPD surveillance of PCV7-vaccinated Navajo or White Mountain Apache native American children at least 18 years old | (Scott et al. 2012; Weinberger et al. 2016) |
| Barcelona | Post-PCV7 | 2007 - 2011 | PCV7-vaccinated healthy children <7 years old in Barcelona | PCV7-vaccinated children with IPD <7 years old in Barcelona | - | (del Amo et al. 2014) |
| France | Post-PCV7 | January 2008 - December 2009 | PCV7-vaccinated healthy children <2 years old across France | PCV7-vaccinated children with IPD <2 years old across France | - | (Varon et al. 2015) |
| Atlanta | Post-PCV7 | June 2008 - August 2009 | PCV7-vaccinated sick Georgia-resident children <5 years old | PCV7-vaccinated children <5 years old with IPD in Atlanta | - | (Sharma et al. 2013) |
| Bogota | Post-PCV7 | June 2011 - November 2011. | PCV7-vaccinated healthy children <18 months old in Bogota | IPD in children <2 years old in Bogota | - | (Parra et al. 2013) |
| France | Post-PCV13 | January 2012 - December 2013 | PCV13-vaccinated healthy children <2 years old across France | PCV13-vaccinated children with IPD <2 years old across France | - | (Varon et al. 2015) |
